# Phenotypic heterogeneity in lag reflects an evolutionarily stable bet-hedging strategy

**DOI:** 10.1101/2025.11.02.686100

**Authors:** Abir George, Ned S. Wingreen, Gautam Reddy

## Abstract

Single-cell experiments in yeast reveal two distinct heritable phenotypes—‘arresters’ and ‘recoverers’—when a clonal population experiences a negative shift in its growth environment. Recoverers exhibit a variable yet finite lag before resuming growth in the new environment, whereas arresters remain in a non-growing, arrested state until more favorable conditions return. Although the diversification of individual cells into arresters and recoverers is a robust phenomenon, it remains unclear whether this coexistence constitutes an evolutionarily stable strategy. Here, we demonstrate that a heterogeneous strategy composed of both arrester and recoverer phenotypes maximizes long-term population fitness across a broad spectrum of growth-lag trade-offs. Our analysis employs a dynamic programming framework to identify the fitness-maximizing distribution of phenotypes for populations that stochastically switch between preferred and non-preferred environments. We propose a minimal model incorporating metabolism, growth, and enzyme allocation to explain the physiological origin of a power-law growth-lag trade-off that favors phenotypic heterogeneity. The theory predicts a nontrivial relationship between the fraction of recoverers and their lag time, which aligns with existing data from wild yeast strains, evolved isolates, and variations in pre-shift growth conditions. This relationship suggests an evolutionary ‘rheostat’-like mechanism that enables populations to rapidly adapt to changing environmental conditions.

## I. INTRODUCTION

Microbes typically live in environments characterized by rapid and unpredictable changes in nutrient availability, temperature, pH, and other critical factors. To persist and thrive under these uncertain conditions, microbes have evolved diverse adaptive strategies. These include mechanisms for physiological adaptation and gene regulation, as well as phenotypic heterogeneity – whereby multiple distinct phenotypes coexist within genetically identical populations. Notably, phenotypic heterogeneity can act as a bet-hedging mechanism, allowing microbial populations to hedge against environmental uncertainty, thus enhancing their long-term success. A well-known example of bet-hedging is the formation of antibiotic-tolerant cells in *Escherichia coli* [1–3]. Within a clonal population, a small fraction of cells grows very slowly, or not at all, which makes them more likely to survive antibiotics and other stressors. These ‘persister’ cells serve as a reservoir for repopulation once favorable conditions are restored. Bet-hedging through phenotypic diversification has been extensively explored in ecological and evolutionary theory, with numerous theoretical studies analyzing its implications [4–12].

Recent experiments in yeast have revealed a distinctive form of phenotypic heterogeneity in response to nutrient shifts. Upon a shift between environments with different nutrients, microbes undergo a period of slow or halted growth, termed the lag phase, during which cells express the machinery that enables growth in the new environment [13]. Yeast (including *Saccharomyces cerevisiae* and its relatives) display a lag period when transferred from a preferred carbon source like glucose to alternative nutrients such as maltose, galactose, or ethanol. Quantitative characterization of this lag phase has revealed considerable variability in the duration and dynamics of lag across natural yeast strains [14–20]: while some strains rapidly resume growth upon a shift, others display many hours of lag before gene expression can reorganize to resume growth.

Beyond this population-level variability observed across strains, single-cell experiments have also found significant heterogeneity in lag. Individual cells from a clonal yeast population display marked variation in lag duration, ranging from immediate resumption of growth to prolonged growth arrest [14, 17]. A standard experimental protocol is to transfer isogenic yeast cells grown, for example, in glucose to maltose and track single-cell growth using time-lapse microscopy (Fig. 1). Cells show two distinct modes of phenotypic response: while some cells (termed ‘recoverers’) resume growth after a few hours, a fraction of cells (termed ‘arresters’) remain in a non-growing, arrested state for the entire duration (~24 hours) of the experiment (Fig. 1b,c). The arrested state is characterized by deformations of subcellular structures [14] and is reversible, that is, arresters resume growth when re-introduced into a glucose-rich environment. While this bimodal response is consistently observed across wild strains and evolved isolates as well as growth media and the time spent in the pre-shift environment, the ratios of recoverers to arresters and the distributions of lag durations within the recoverer sub-population both vary substantially across these scenarios [17, 20]. A natural question is whether this coexistence of arrester and recoverer states reflects a stable evolutionary strategy.

**FIG. 1.**
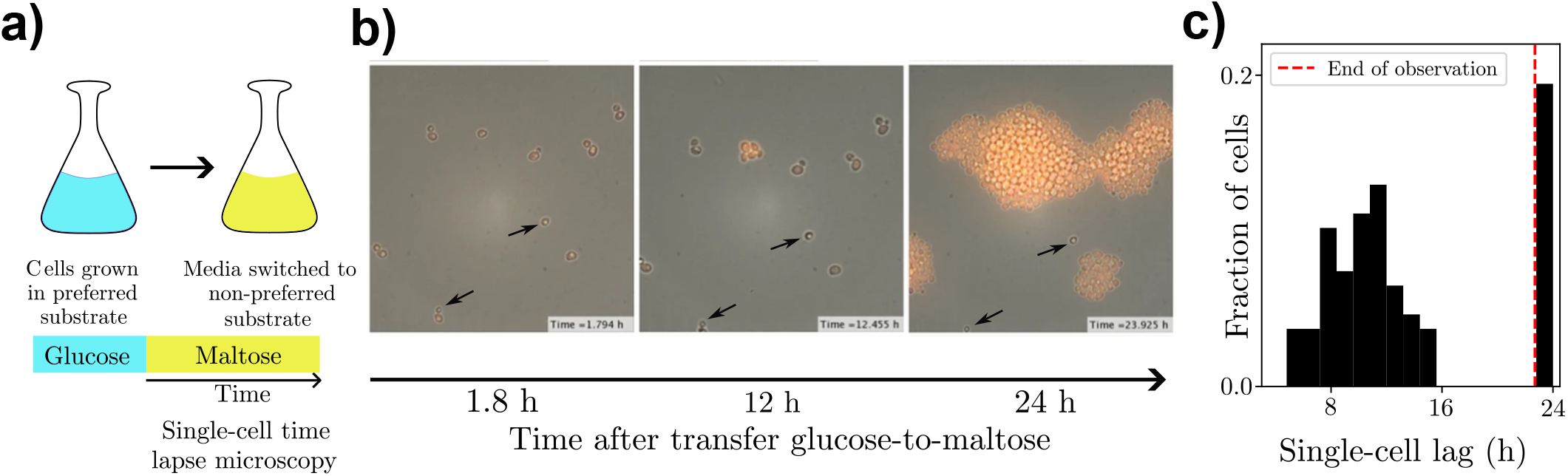
Heterogeneity in single-cell lag durations following a glucose-to-maltose shift. (a) Schematic of experimental protocol: yeast cells are pre-grown in glucose, then the media is switched to maltose where cells are tracked using time-lapse microscopy. (b) Time-lapse images of a yeast population following a sudden switch from glucose to maltose. MAL12-yECitrine fluorescence indicates growth resumption. Images taken at 1.8 h, 12 h, and 24 h after the switch. Cells exhibit heterogeneous behavior in lag phase recovery, including a subpopulation that remains arrested (indicated by black arrows). Images reproduced from [17]. (c) Distribution of single-cell lag durations across the population. The red dashed line marks the end of the observation period (23.5 h), beyond which lag durations were not measured.

If rapid recovery from lag entailed a negligible physiological cost (for example, in pre-shift growth rate), then we should expect cells to prefer a homogeneous strategy that recovers as quickly as possible. However, the observation of an arrested state suggests that cells face intrinsic limitations when adapting to maltose or other less-preferred carbon sources. Extensive experimental work has characterized trade-offs between growth and adaptability in microbes [14, 16, 18, 21–26]. When such trade-offs are present, the coexistence of distinct phenotypes can emerge as an evolutionarily stable strategy [14, 23–25, 27]. Thus, one possible explanation for the observed coexistence is that individual yeast cells are constrained by some nontrivial physiological trade-off that prevents them from simultaneously achieving rapid preshift growth and rapid post-shift recovery. This hypothesis is supported by measurements which show that arresters grow faster than recoverers in the pre-shift environment [14].

To determine the evolutionarily stable strategy in the presence of such a growth-lag trade-off, we leverage a novel optimization framework based on dynamic programming. The framework leads to a general decision-making principle that can be used to find fitness-maximizing bet-hedging strategies for a population fluctuating between a preferred and a non-preferred environment. We show that, depending on environmental statistics and the physiological trade-off, the population either adopts a bet-hedging strategy composed of two “specialist” phenotypes (arresters and recoverers) that respectively are well-suited for growth and rapid recovery, or picks a “generalist” phenotype that balances both traits. We propose a minimal metabolic model that explains the physiological origin of a power-law-like growth-lag trade-off. Furthermore, the model predicts that populations experiencing different environmental statistics follow a characteristic relationship between the ratio of recoverers and arresters, and the lag duration of recoverers. We test this prediction using existing experimental data, which hints at an underlying evolutionary mechanism that enables rapid adaptation to changes in environmental statistics.

## II. THEORETICAL FRAMEWORK

To quantify the long-term fitness benefits of phenotypic heterogeneity, we developed a cellular decision-making framework based on dynamic programming [28]. We first describe the general framework before specializing to the growth-lag scenario. Another application of this framework to a growth-death scenario is presented in Appendix B 4.

Consider a microbial population where cells adopt one of *K* phenotypic states, labeled by *k*, each of which is characterized by its growth rate *γ*(*s, k*) in state *s*. The state *s* can represent, for example, the time since a nutrient switch, the concentration of a nutrient, or any history-dependent quantity tracked by the cell. The instantaneous growth rate of a population of *N* cells at time *t* is

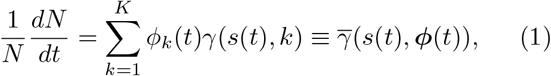

where *ϕ*_*k*_ is the fraction of cells with phenotype *k* and ***ϕ*** = (*ϕ*_1_, *ϕ*_2_, …, *ϕ*_*K*_) is the phenotype distribution. The long-term growth rate Γ of the population is then

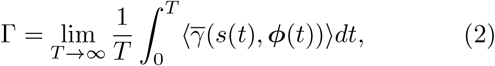

where the angled brackets denote an expectation over changes in state.

We focus on populations that transition repeatedly between preferred (P) and non-preferred (NP) environments, such as yeast shifting between glucose and maltose. Importantly, we assume that in P the population can rapidly reorganize gene expression and thereby adjust its phenotype distribution, whereas cells remain phenotypically locked upon the switch to NP. This environmental switching model is schematically illustrated in Fig. 2a,b. Fluctuating environments will select for phenotype distributions that best balance overall population growth in P and NP. A key conceptual question is when and whether this best population-level strategy consists of one, two, or multiple phenotypes (Fig. 2c).

**FIG. 2.**
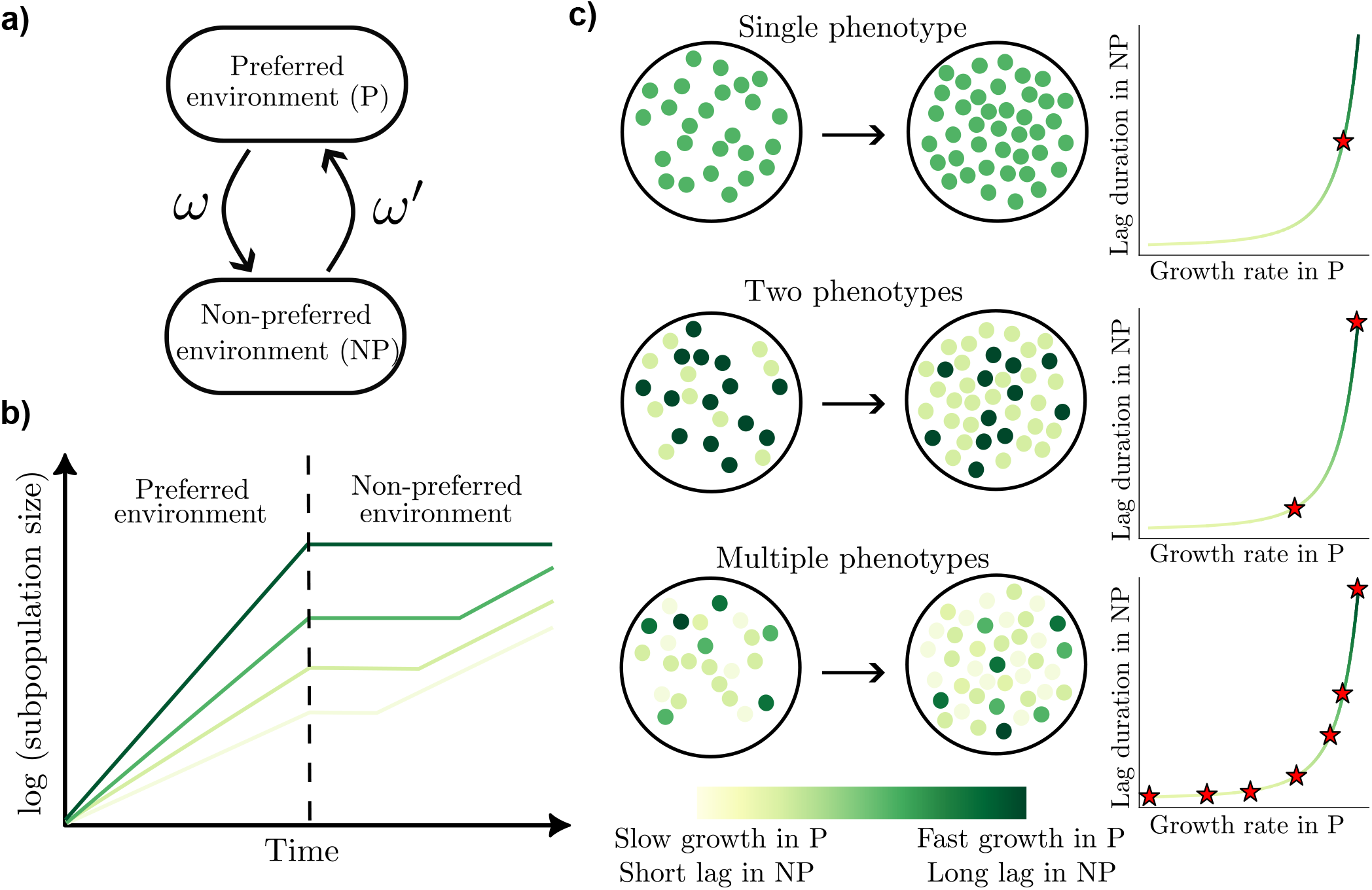
Phenotypic strategies in fluctuating environments. (a) Model of environmental switching with transitions between a preferred (P) and a non-preferred (NP) environment occurring at rates *ω* and *ω*′, respectively. (b) Growth dynamics of pure phenotype subpopulations in a fluctuating environment. Distinct phenotypes grow at different rates in the preferred environment (P) and experience varying lag durations upon transition to the non-preferred environment (NP), with a trade-off between growth rate in P and lag duration in NP. Cells that grow slowly in P recover more quickly and resume growth earlier in NP. As a result, these phenotypes increase in relative abundance during NP. (c) Schematic showing possible strategies for a population of cells: a single phenotype (top), two phenotypes (middle), or multiple phenotypes (bottom). Disks represent individual cells colored by their phenotype, corresponding to a position on the growth-lag trade-off curve (right). Red stars indicate the phenotypes that comprise the strategy.

Our objective is thus to identify the optimal phenotype distribution ***ϕ***\* in P that maximizes the long-term growth rate Γ in the face of environmental fluctuations. Given state *s*, and our assumption of rapid phenotype adjustment in P, we show that

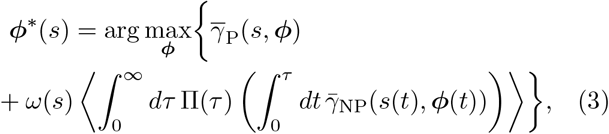

where Π(*τ*) is the probability density of the duration *τ* spent in NP before switching back to P, *ω*(*s*) is the instantaneous rate of switching from P to NP and we have included the subscripts P and NP to 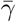 for clarity. Equation 3 has an intuitive interpretation: the optimal strategy balances the instantaneous growth rate in P (first term) and the expected long-term growth rate if the environment were to transition to NP, scaled by the probability that the transition occurs (second term).

## III. RESULTS

### Optimal strategies given a growth-lag trade-off

In our growth-lag scenario, cells with phenotype *k* have a growth rate *γ*_*k*_ in P and a lag duration *ℓ*_*k*_ in NP with no growth during lag. After recovering from lag, cells grow in NP at a common rate *γ*′. Phenotypes experience a growth-lag trade-off: phenotypes that grow faster (slower) in P exhibit longer (shorter) lag durations in NP (Fig. 2b). To incorporate the effect of lag, the state *s*(*t*) in NP tracks the time since the most recent transition from P to NP. For simplicity, we drop any dependence of ***ϕ*** on *s* in P, noting however that the results also apply to scenarios where ***ϕ***\* depends on the nutrient in P or historical information that could be used to anticipate a switch. The optimization objective then becomes

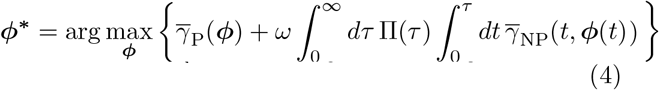

from which we derive the specific optimization objective

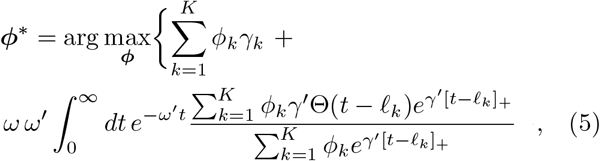

where *ω*′ is an assumed constant rate of switching from NP back to P so that 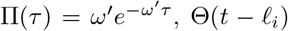 is the Heaviside function and [*t* − *ℓ*_*i*_]_+_ = max(0, *t* − *ℓ*_*i*_). Integrating Eq. 5 by parts, the objective function can be simplified to

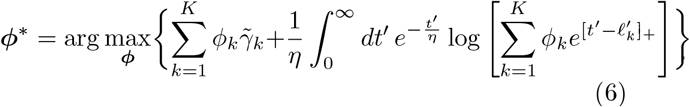

where 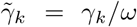 and 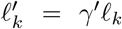 represent normalized growth rates and lag durations respectively, and *η* = *γ*′*/ω*′ is the average number of generations of a no-lag subpopulation in NP.

A key factor determining a population’s optimal strategy in a fluctuating environment is the form of the growth-lag trade-off. To capture the range of possible behaviors, we consider a family of growth-lag trade-offs:

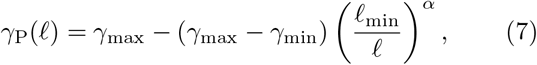

where lag *ℓ* ≥ *ℓ*_min_, and the growth rate in P lies in the range *γ*_min_ ≤ *γ*_P_ ≤ *γ*_max_. The parameter *α* defines a family of trade-off curves, with larger *α* corresponding to larger *γ*_P_ for the same lag *ℓ*. This parametric form always includes phenotypes that have maximum growth in P at the cost of infinite lag (*ℓ* = ∞) in NP.

### Numerical result: at most two phenotypes, with one arrester

We numerically computed the optimal phenotype distribution over a continuous set of phenotypes constrained by the above growth–lag trade-off curve (Eq. 7). Although one might have expected the optimal strategy to exploit a broad range of phenotypes, we found that the optimal distribution contains at most two distinct phenotypes. This restriction to no more than two phenotypes applies across the whole parameter space defined by the environmental transition rates (*ω, ω*′), as illustrated in Fig. 3a,b.

**FIG. 3.**
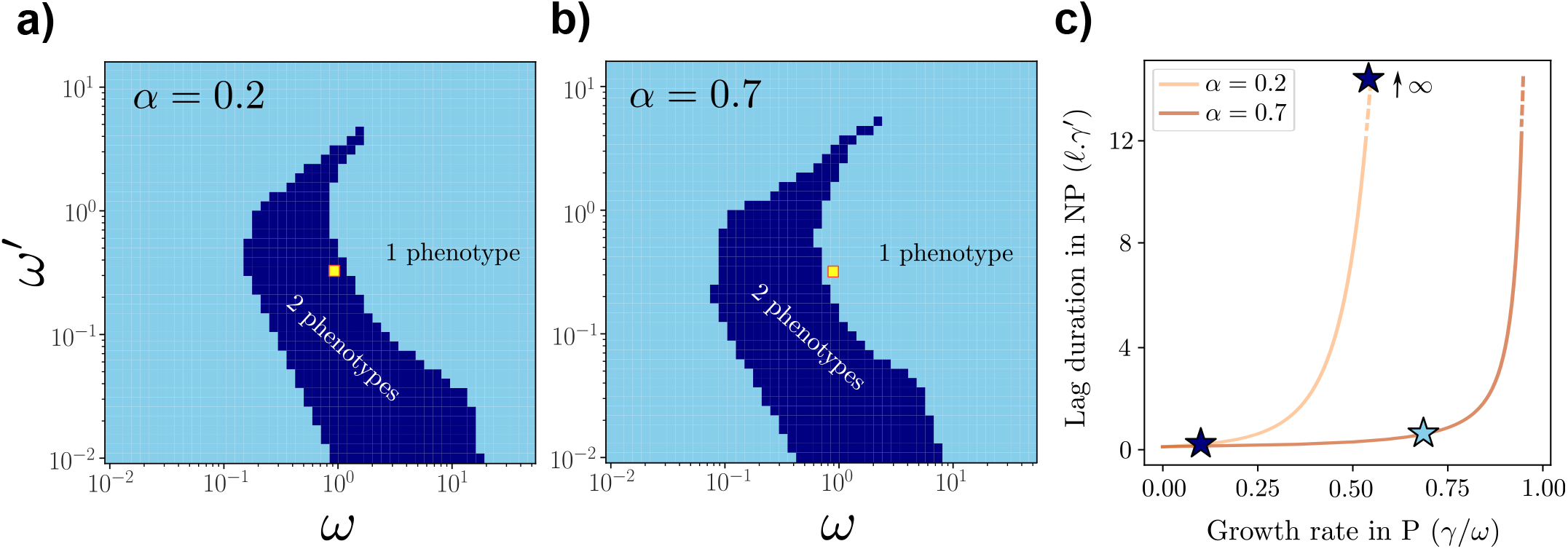
Optimal phenotypic strategies consist of at most two distinct phenotypes. (a, b) Phase diagrams of the optimal number of phenotypes over the space of environmental switching rates, *ω* and *ω*′, for fixed trade-off curves parameterized by *α* = 0.2 (a) and *α* = 0.7 (b). Light blue regions correspond to a single optimal “generalist” phenotype, while dark blue regions correspond to two coexisting “specialists”, an arrester and a recoverer. (c) Different trade-off curves between growth rate in P and lag duration in NP from A and B, with optimal phenotypes for the (*ω, ω*′) values indicated by yellow squares. For increasing *α*, the optimal strategy transitions from a two-phenotype (specialist) solution (dark blue stars) to a single-phenotype (generalist) solution (light blue stars).

Within the two-phenotype regime, the optimal strategy consistently comprises one phenotype with finite lag – a recoverer – and another phenotype with infinite lag (*ℓ* = ∞) – an arrester. The recoverer is adapted to resume growth in the non-preferred environment (NP), while the arrester is specialized for maximal growth in the preferred environment (P). The optimal strategy assigns a nonzero probability to the arrester phenotype, even though it never grows in NP. This is perhaps a surprising result: it implies that under certain environmental conditions, it is beneficial for a population to maintain a subpopulation that entirely forfeits the ability to grow in one environment in order to maximize performance in the other, even though both environments can support growth. Similarly, the recoverer phenotype sacrifices growth in P in order to perform better in NP. The two phenotypes in this bet-hedging strategy can thus be viewed as *specialists*, each appropriately suited for a distinct environmental state.

It is clear that the phase diagram is re-entrant, that is, for increasing *ω/ω*′, the optimal strategy goes from generalist to specialist, and back to generalist. For small *ω/ω*′, the generalist strategy corresponds to having only the arrester in the population since the environment is mostly P. By contrast, for *ω/ω*′ ≈ 1, the population should optim, ally adopt a specialist strategy (with deviations with respect to this ratio for *ω, ω*′ ≪ 1 and *ω, ω*′ ≫ 1). When *ω/ω*′ increases beyond this, the optimal strategy reverts to a generalist strategy with a recoverer phenotype. This makes sense as the population spends more time in NP and the recoverer phenotype is able to maximally contribute to the population growth.

As shown in Fig. 3c, varying the shape of the growth–lag trade-off via the power-law exponent *α* can qualitatively change the optimal strategy for fixed (*ω, ω*′). For example, for the (*ω, ω*′) values in the yellow squares in Fig. 3a,b, at low *α* the optimal strategy is two specialists (one being an arrester), while at higher *α* the optimal strategy is a single *generalist* phenotype that balances growth in P and lag in NP.

### Phenotypic distributions with a few discrete phenotypes are optimal

The observation that the optimal phenotype distribution contains a few (here, at most two) discrete phenotypes can be explained in the special case when the time spent in the non-preferred environment (NP) is fixed at a constant value *t*. In this case, the optimization objective in Eq. 6 reduces to (with *t*′ = *γ*′*t*)

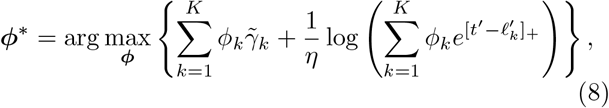

with the constraints 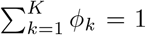 and *ϕ*_*k*_ ≥ 0 for all *k*. The Karush-Kuhn-Tucker (KKT) theorem provides a set of conditions satisfied by ***ϕ***\* subject to these constraints. The KKT conditions imply that the growth rates and lag durations of any pair of phenotypes (indexed by *m* and with nonzero probability are related according to

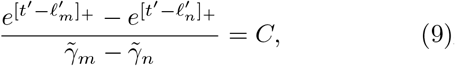

for some constant *C*. This restrictive condition may be true for a continuous set of phenotypes in certain special scenarios, for example, when 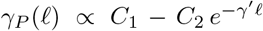, but that will not be true in most scenarios, including for the family of growth-lag trade-offs in Eq. 7. Some geometric intuition can be obtained by noting that solutions ***ϕ***\* for an objective of the form Eq. 8 will lie on the vertices of a region formed by the intersection of a hyperplane with normal vector ***γ*** = (*γ*_1_, *γ*_2_, …, *γ*_*K*_) and the probability simplex. These vertices are points with nonzero probability for at most two phenotypes. For complex dwell time distributions in NP, it is possible that the optimal phenotypic distribution has more than two phenotypes. However, we consistently find at most two phenotypes for the exponentially distributed dwell times considered in our numerical simulations, presumably because this distribution has a single typical timescale.

### The geometry of the growth-lag trade-off determines whether a generalist or specialist strategy is preferred

When is a single generalist phenotype favored over a mixture of two specialist phenotypes? Optimizing Eq. 4 for the single best phenotype solution, the lag *ℓ*_g_ of the generalist phenotype (provided it is finite) satisfies 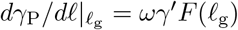, where 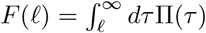 for general growth-lag trade-offs *γ*_P_ and dwell time distributions Π(*τ*) (Appendix D). Considering phenotypes in the neighborhood of *ℓ*_g_, we find that a mixed strategy is preferred if (Appendix D)

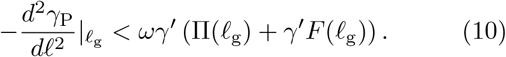

This condition implies that the curvature of the growth-lag trade-off around the generalist solution determines whether a pure or a mixed strategy is favorable. Moreover, the solutions on the two sides of the transition point (i.e., when equality holds in Eq. 10) are discontinuous: extreme specialists are preferred whenever a mixed strategy is optimal (Appendix D).

### The optimality of the arrester phenotype

A consistent observation in our numerical simulations is the selection of a perfect arrester phenotype (with infinite lag) whenever a mixed strategy is preferred. The selection of this arrester phenotype can be explained by a heuristic argument based on how the growth-lag trade-off scales at long lags and the tail probabilities of dwell times in NP (see Appendix E 1 for a longer discussion). Since the distribution of dwell times in NP decays exponentially, the marginal cost of increasing the lag of the arrester phenotype also falls off at the same exponential rate. On the other hand, the marginal benefit in pre-shift growth rate declines more slowly according to a power-law given by Eq. 7. Thus, for long lags, any increase in lag of the arrester delivers a net positive marginal benefit, implying that it is always beneficial to increase the lag indefinitely.

### A minimal metabolic model reproduces a powerlaw-like trade-off between pre-shift growth rate and post-shift lag duration

How might a power-law-like growth-lag trade-off arise? In a model where different nutrients are metabolized by independent pathways, it is not clear why a strong growth-lag trade-off should pertain to a nutrient switch. The allocation of catalytic enzymes for the two pathways could be independent, allowing growth and lag to be decoupled and separately optimized. However, studies in bacteria [21, 22] reveal that cells primed to grow quickly on a glycolytic nutrient experience a lag upon switching to a gluconeogenic nutrient. In particular, Basan et al. [21] report a reciprocal relationship between the steady-state growth rate in different glycolytic substrates and the corresponding adaptation times after a shift to acetate. Schink et al. [22] provide a mechanistic interpretation associated with flux reversal in the glycolysis and gluconeogenesis pathways. Guided by these observations, we developed a minimal metabolic model (Fig. 5) that results in a power-law-like trade-off between pre-shift growth rates and post-shift lag durations.

In our model, two nutrients are processed along the same pathway, but the metabolic flux is controlled by catalytic enzymes that favor opposing directions (Fig. 4a). A key feature of this model is that after a nutrient switch, the imbalance in enzyme abundances which enabled flux in one direction must be reversed to allow flux in the other. This reversal in turn requires biomass production, which is slow until flux can be reversed. The coupling between flux reversal and biomass production consequently leads to a long lag. Thus, this minimal model is sufficient to obtain a growth-lag trade-off.

**FIG. 4.**
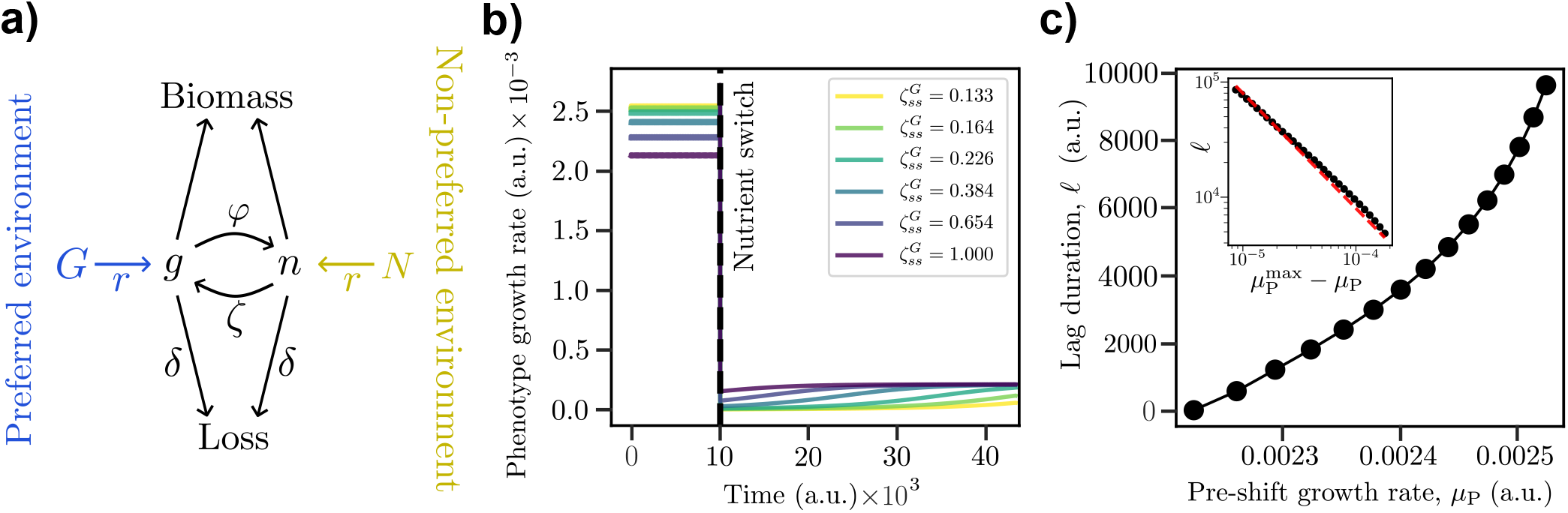
A minimal metabolic model with a growth-lag trade-off. (a) Schematic showing two central metabolites *g* and *n* where enzymes *ϕ* catalyze *g* → *n*, and enzymes *ζ* catalyze *n* → *g*. In the preferred environment, nutrient *G* is present and *N* is absent. Allocating enzymes to high *ϕ* achieves a high growth rate in this environment. However, after a switch to the non-preferred environment, pre-existing high *ϕ* prolongs the reaction *g* → *n*, which delays net flux reversal, producing a lag in growth until *ϕ* falls and *ζ* rises. (b) Time courses around the nutrient switch (vertical dashed line) showing the growth rates of different enzyme-allocation phenotypes. We define “lag” to end when the post-shift growth rate exceeds half the steady-state growth rate *µ*_NP_ in the non-preferred environment. (c) Growth–lag trade-off curve generated by varying pre-shift enzyme allocation, keeping 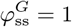 fixed while decreasing 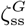 from 1 to 10^−1.8^. This monotonically increases pre-shift growth rate *µ*_P_ but prolongs post-shift lag *ℓ*. The inset shows the predicted power-law scaling with exponent − 1 (red dashed line) for *ν* = 2. Details of the model are given in Appendix G and the parameter values are listed in Table I.

**FIG. 5.**
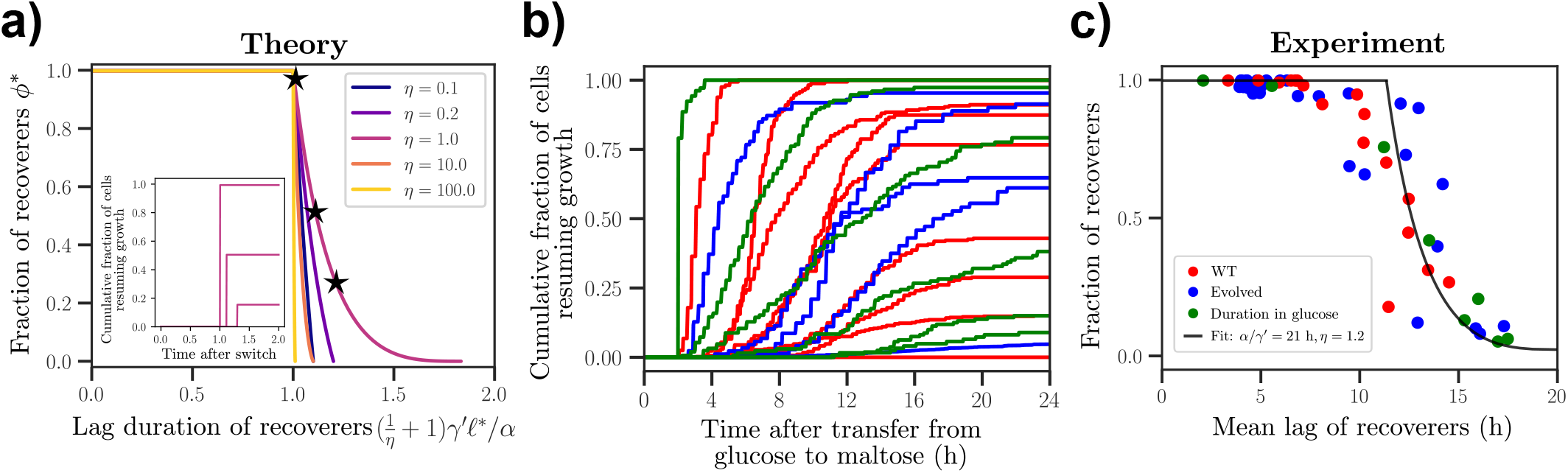
Comparison of theoretical predictions and experimental data for fraction of recoverers versus lag duration. (a) Theoretical relationship between the optimal fraction of recoverers and their lag duration for different values of *η* = *γ*′*/ω*′. The lag duration is rescaled using parameters *η, γ*′ and *α*. Inset – Theoretical curves showing the single-cell lag duration after a switch from the preferred environment to the non-preferred environment. Different curves correspond to different points (black stars) on curve for *η* = 1. Units for the x-axis match that of the main figure. (b) Experimental measurements of yeast single-cell lag duration after a switch from glucose to maltose, recorded via time-lapse microscopy. Each curve corresponds to the cumulative fraction of cells resuming growth after the switch. Red curves represent measurements for different wild strains, blue curves represent measurements for evolved isolates that experienced repeated cycling of glucose-maltose media, and green curves correspond to populations that experienced different pre-shift durations in glucose. Only a subset of these traces is shown here due to limited space. See Fig. S2 for more. Data from Ref. [15, 16]. (c) Experimental plot of the fraction of recoverers against the mean lag duration of the recoverers. Red, blue, and green dots represent, respectively, wild strains, evolved isolates, and variations based on pre-shift durations in glucose. The solid black curve is a fit to our theoretical prediction (Eq. 11) relating the recoverer fraction to the lag duration, yielding *α/γ*′ = 21 h and *η* = 1.2.

**TABLE I.** Values for parameters used in the metabolic model.

| Parameters | Values |
| --- | --- |
| $G_P, N_P$ | 2.0, 0.0 a.u. |
| $G_{NP}, N_{NP}$ | 0.0, 1.0 a.u. |
| $r, \delta$ | 1.0, 3.0 a.u. of 1/time |
| $K_g, K_n, K_{BM}$ | 1.25, 2.0, 0.75 a.u. |
| $\nu$ | 2 |

We quantify growth in the preferred environment (P) by the steady-state growth rate *µ*_P_ and define lag, *ℓ*, in the non-preferred environment (NP) as the time for resumed growth to exceed half the NP steady-state growth rate *µ*_NP_ (i.e. *ℓ* is the first-passage time for post-shift growth rate ≥*µ*_NP_*/*2). The model tracks two central metabolites, *g* and *n*, interconverted by two opposing enzyme activities: *ϕ* catalyzes *g* → *n* and *ζ* catalyzes *n* → *g* as shown in Fig. 4a. Metabolic fluxes are Michaelis–Menten, growth rate via biomass production *µ* is a cooperative function of *g* and *n*, and enzyme levels relax toward environment-specific set points via

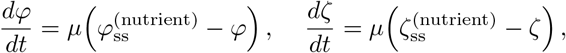

where the instantaneous growth rate is 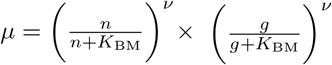 and 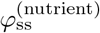 and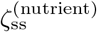 are the steadystate concentrations of enzymes *ϕ* and *ζ* for a specified nutrient, respectively. For simplicity, the system is taken to be symmetric with respect to the two metabolites, except for the externally supplied nutrients. The rate constant *r* governs the influx into metabolites *g* and *n*, contributing *rG* and *rN* from *G* and *N*, respectively. We initialize the system at steady state in P with *G* at a fixed non-zero concentration *G*_P_ and *N* = 0, then impose an instantaneous environmental switch to NP by changing nutrient availability so that *G* = 0 and *N* has non-zero concentration *N*_NP_ *< G*_P_.

After the nutrient switch, high pre-existing *ϕ* suppresses the now-desirable net flux from *n* → *g* and leads to excess accumulation of *n*. This depletes cellular resources and delays the reversal of net flux. Slow biomass production gradually rebalances the enzymes, decreasing *ϕ* and increasing *ζ*, allowing the growth rate *µ* to rise to its steady-state value in NP. This rise time is an effective post-shift lag.

For each value of 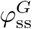, we vary 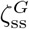 to obtain a growthlag trade-off. The values of 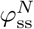 and 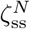 are kept fixed. As described below, we find that the metabolic model operates in at least three qualitatively distinct regimes for different values of 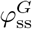 depending on how the system produces new enzymes *ζ* after the switch from P to NP. Importantly, we obtain a power-law growth-lag trade-off for intermediate values of 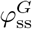. In Fig. 4b, we show specific time courses of the growth rate for different enzyme allocations (recoverer-like versus arrester-like) in this in-termediate regime of 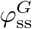.

First, when 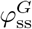 is sufficiently small (and pre-shift growth is minimal), the depletion of *g* is gradual such that there is some growth (and thus production of *ζ*) for a short period after the switch from P to NP. The brief production of *ζ* is sufficient to avoid a prolonged lag phase as any non-zero *ζ* helps reverse the flux from *n* → *g*, promoting growth and subsequent production of *ζ*. That is, the system avoids a lag phase at the expense of minimal pre-shift growth.

Second, when 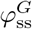 is sufficiently large, the value of 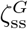 that maximizes growth in P is non-zero (Fig. S5). This is because a large 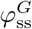 rapidly depletes *g* even with influx from *G* in P. It is beneficial to oppose some of the flux from *g* → *n* via *ζ* to maintain a pool of *g* and *n* and promote growth. The pre-shift growth rate *µ*_P_ decreases quadratically with small deviations of 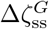 from the growth-optimal allocation, that is, 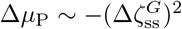. Since 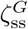 is non-zero, the post-shift lag *ℓ* is finite even when pre-shift growth *µ*_P_ is maximal and *ℓ* decreases lin-early with increasing 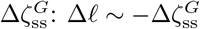. These two relations then lead to a quadratic relationship Δ*µ*_P_ ~ − Δ*ℓ*^2^ near the growth-optimal 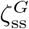 for a given 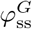. While the lag at maximal growth is finite, maintaining a large pool of enzymes 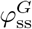 and 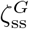 would incur a significant energetic cost. That is, the system could optimize pre-shift growth and still have finite lag after the switch but only by paying the energetic and resource cost of simultaneous expression of *ϕ* and *ζ*, which would lead to futile catalytic cycles [22].

Third, for intermediate values of 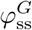, the growth-optimal allocation is 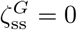 (Fig. S5). The rapid depletion of *g* after the switch and the lack of any conversion from *n* → *g* leads to infinite lag when 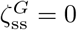. Any small, non-zero 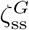 decreases growth rate linearly, 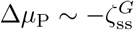. Importantly, we show analytically that 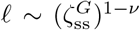 to first order in 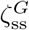 (for *ν >* 1). This relation arises from the slow production of *ζ* after the switch. The two relationships, 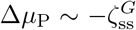 and 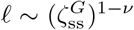, together imply a power-law-like growth-lag trade-off Δ*µ*_P_ ~ − *ℓ*^−*α*^ with exponent *α* = (*ν* − 1)^−1^ (see Appendix G for more details). This predicted power-law relationship was verified in simulations (Fig. 4c).

### Theory predicts a relationship between the fraction of recoverers and their lag

Our theory implies that for every choice of environmental statistics (specified by *ω* and *ω*′ in our model), there is a unique pheno-type distribution that maximizes long-term growth. In other words, optimality requires that different features of the phenotype distribution (such as the fraction of recoverers and their lag) are not independent but rather co-vary with variations in environmental statistics. To derive these constraints, we now consider a simplified model where the phenotype distribution is restricted to at most two phenotypes. In this model, the phenotype distribution is parameterized by two quantities: the lag of the recoverer (denoted *ℓ*) and the fraction of recoverers (denoted *ϕ*). The remaining fraction 1 − *ϕ* of arresters has infinite lag. For such a two-phenotype system, cells can select *ϕ* in P by stochastically switching between recoverer and arrester phenotypes. The difference in pre-shift growth rates between the arrester and recoverer phenotypes and the switching rates fix *ϕ* at steady-state (Appendix F).

Optimality of the phenotype distribution predicts that *ℓ* and *ϕ* co-vary according to a nontrivial relationship across different environmental statistics. For exponential dwell times in NP and the growth-lag trade-off in Eq. 7, we derived an analytical relationship between the optimal fraction *ϕ*\* of recoverers and their optimal lag *ℓ*\* (for details see Appendix E 2). Whenever 0 *< ϕ*\* *<* 1, the relationship is given by

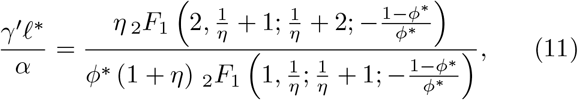

where _2_*F*_1_ is a hypergeometric function and recall that *η* = *γ*′*/ω*′. As *ϕ*\* → 1, we have *γ*′*ℓ*\**/α* = *η/*(1 + *η*) and as *ϕ*\* → 0, *γ*′*ℓ*\**/α* = min {1, *η*}, which delineate the two boundaries of the coexistence region.

Figure 5a illustrates the predicted relationship between the optimal fraction of recoverers (*ϕ*\*) and their optimal lag (*ℓ*\*) for different values of *η*. We obtain a family of curves that show a consistent inverse relationship: *ℓ*\* initially increases while maintaining a single recoverer population (*ϕ*\* = 1) with decreasing switching rate *ω* from P to NP. For a sufficiently small value of *ω*, the population splits into two phenotypes (0 *< ϕ*\* *<* 1) while the lag of the recoverer continues to grow until the population again consists of a single phenotype (*ϕ*\* = 0). The ratio between the values of *ℓ*\* at the two boundaries of the coexistence region, *ϕ*\* → 0 and *ϕ*\* → 1, is at most two (achieved when *η* = 1), imposing a constraint on the possible shapes in this family of curves.

An important point is that the relationship between the rescaled lag (in units of *α/γ*′) and fraction of recoverers is only mediated by the parameter *η* (the expected number of generations in NP of a no-lag phenotype). It is agnostic to factors such as genetic variability among different strains and dwell time distributions in the preferred environment, making it broadly applicable across diverse experimental conditions and yeast strains. We can thus quantitatively test this prediction for various environmental conditions and genetically distinct yeast populations using existing datasets.

### Experimental evidence for a relationship between fraction of recoverers and their lag

We analyzed experimental data from single-cell measurements of yeast populations undergoing a glucose-to-maltose transition (Fig. 5b). These data report the cumulative fraction of cells resuming growth after a switch from media containing glucose to media containing maltose for up to 24 hours after the switch [16]. In Fig. 5b, each individual red curve represents lag measurements for a different wild strain of yeast, blue curves represent measurements taken for evolved isolates of a long-lag strain in parallel cultures after experiencing 6-8 cycles of alternating glucose and maltose-containing media, and green curves represent measurements for populations that experienced different glucose pre-growth durations. From the data, we obtained distributions of lag durations for each tracked population.

A consistent feature is the bimodality of these distributions, delineating two distinct subpopulations of recoverers, which resumed growth before the experiment ended, and arresters, which remain arrested. We calculated the average lag duration of the recoverers and their corresponding fraction *ϕ* within each population. We observed a robust inverse relationship where populations with longer lags had smaller fractions of recoverers (Fig. 5c). This empirical trend matches well with predictions from our two-phenotype model. Treating *η* and *α/γ*′ as fitting parameters, we obtained best-fit values of *η* = 1.2 and *α/γ*′ = 21 h. Using the experimental estimate *γ*′ = 0.1 h^−1^ from [16], these values correspond to *α* = 2.1 and *ω*′ = 0.08 h^−1^. See Fig. S4 for the phase diagram for the optimal number of phenotypes with these physiological and environmental parameters.

## IV. DISCUSSION

Microbial populations repeatedly exposed to environmental shifts must reconcile fast growth in the current niche with the capacity to resume growth after a switch. In this study, we investigated the role of phenotypic heterogeneity in microbial populations, focusing on the adaptation of yeast to changing carbon sources. The central experimental observation motivating this work is that following a switch from a glucose-rich environment to a maltose-rich environment (or other environments), a substantial fraction of genetically identical yeast cells remain arrested in lag phase for the entire duration of the experiment [14–17]. This persistent growth arrest suggests that cells in this subpopulation occupy a distinct physiological state, raising the question whether this heterogeneity might reflect an evolved strategy to optimize population-level fitness in fluctuating environments. To address this, we developed a general framework based on dynamic programming to determine the optimal bet-hedging strategies for microbial populations transitioning between environments where cells can vs. cannot readily reorganize gene expression. We found that the optimal phenotype distribution consists of at most two discrete phenotypes, even when a continuous range of phenotypes is allowed. Under certain conditions, the population adopts a bet-hedging strategy composed of two distinct specialists: one phenotype minimizes lag to recover quickly in the non-preferred environment (NP), while the other maximizes growth in the preferred environment (P) but never recovers in NP. In other regimes, a single optimal generalist phenotype emerges that balances growth and lag.

A key prediction from our model is a nontrivial relationship between the fraction of recoverers and their lag. This relationship is a consequence of imposing optimality of the phenotype distribution for a given physiological trade-off across different environmental transition rates. This prediction matches single-cell datasets spanning wild strains, evolved lines, and varying pre-growth nutrient durations, suggesting an underlying constraint in the genetic circuit that mediates phenotypic diversification.

Two assumptions underpin our results that populations benefit from having phenotypic heterogeneity with distinct discrete phenotypes: First, the population can re-tune its phenotype distribution rapidly in the preferred environment (P) but phenotypes are locked during lag in the non-preferred environment (NP). Second, a monotonic trade-off links higher pre-shift growth to longer post-shift lag. Other ingredients of the model, such as exponentially distributed dwell times, the exact form of the trade-off, and the perfect arrester limit (*ℓ* → ∞), allow analytic tractability but are not required for heterogeneity to be favored. More complex dwell-time distributions could create regimes where more than two distinct specialists make up the optimal strategy, each tuned to a different characteristic timescale of environmental shifts. We note that single-cell lag variability is not always strongly bimodal. In some bacterial contexts, lag times can span wide distributions, sometimes with long tails. For example, *Escherichia coli* can exhibit broad lag-time distributions following nutrient transitions or starvation, with growth resumption dominated by short-lag sub-populations, while long-lag cells can contribute to stress tolerance [29, 30]. Our prediction of at most two discrete phenotypes should be interpreted as a statement about the optimal strategy within our framework’s assumptions, namely, that there are two types of environments (P and NP) and that the distribution of times spent in the NP environment has a single timescale. Relaxing either assumption could lead to a broader optimal phenotype distribution.

Mechanistically, a minimal metabolic model motivates why a trade-off may exist: allocating enzymes to drive flux in the pre-shift direction (fast growth) delays net flux reversal and consequently delays resumed growth after the switch. Systematic variation of pre-shift enzyme allocation reproduces a power-law growth-lag trade-off curve consistent with the phenomenology encapsulated by Eq. 7. Prior work has also shown that growth-lag trade-offs can arise from constraints on flux reversal and proteome allocation in central metabolism [21, 22], though the variation in pre-shift growth rates in these cases is due to different pre-shift glycolytic substrates rather than heterogeneous growth rates on a single substrate.

Our framework extends classical bet-hedging theory [3, 6, 7, 12] in a key way: We couple pre-shift growth and post-shift lag through an explicit metabolic constraint that encodes how proteome allocation before the switch creates a bottleneck in the speed of adaptation after the switch. This makes the trade-off mechanistic rather than just phenomenological. Our work is thus novel in that we link cellular physiology to population level growth outcomes in fluctuating conditions.

Lag heterogeneity following carbon shifts in *S. cerevisiae* shares a key similarity with persister formation in *E. coli*, competence in *Bacillus subtilis*, and sporulation in filamentous fungi: all involve a slow-growing subpopulation that sacrifices immediate proliferation for future advantage. In these systems, phenotypic diversity provides a benefit to the population by being prepared for unfavorable conditions. The “recoverer” state in our model plays this protective role, maintaining the population (e.g. against competitors) until favorable conditions return.

Our analysis suggests that the recoverer lag-fraction relationship can function as an evolutionary “rheostat”, providing populations with a mechanism to tune their balance between fast growth and lag. In our metabolic model, such a rheostat could be wired through regulatory control of proteome allocation. Changes in the expression of enzymes that bias flux toward pre-shift growth would continuously shift both the growth rate in the preferred environment and the lag after a nutrient switch. This would enable populations to smoothly adjust their phenotypic distribution in response to the statistics of environmental change, without requiring rewiring of their metabolic circuitry. More broadly any modulation (genetic or otherwise) of upstream regulatory pathways that control the phenotype distribution will then satisfy an optimality constraint that can be tuned to the rate of environmental change, offering a convenient evolutionary rheostat for rapidly adapting to new environments. In short, with a single control knob, closely related strains constrained by the same trade-off could flexibly adapt to different environmental fluctuation statistics.

Promising directions for future experimental work include (i) varying *ω* and *ω*′ (e.g. in a microfluidic setup) to test the predicted re-entrant generalist–specialist–generalist behavior, (ii) adjusting pre-shift enzyme allocation (genetically or via growth history) to test the coupling between growth and lag, and (iii) measuring the survival of long-lag arresters to test for a “perfect arrester” state. Together, these experiments would show whether the recoverer fraction–lag relation functions as an evolutionary rheostat across strains.

## DATA AND CODE AVAILABILITY

The data and code used to generate all figures are available at https://doi.org/10.5281/zenodo.17582890.

## ACKNOWLEDGMENTS

We thank Piyush Nanda and Andrew Murray for many fruitful discussions. This work was supported in part by Princeton University through the Center for the Physics of Biological Function, and in part by the National Institutes of Health under award number R01GM082938 (NSW). GR was partially supported by a joint research agreement between NTT Research Inc. and Princeton University. The content is solely the responsibility of the authors and does not necessarily represent the official views of the National Institutes of Health.

## Appendix A: Experimental data

Data for single-cell lag phase measurements was taken from New et al. 2014 and Cerulus et al. 2018. In short, cells were pre-grown in glucose, harvested, and transferred to media containing maltose. The lag duration of individual cells was measured using automated timelapse microscopy, defined as the time from transfer until the first observable sign of division (bud emergence or resumption of bud growth). For experimental details, refer to New et al. 2014. To generate the distribution of single-cell lag durations (Fig. 1b), data was specifically taken from Dataset S1. It contains single-cell lag phase measurements for 18 strains of *S. cerevisiae* and we picked the distribution for strain ‘DBVPG1373’ for Fig. 1b in the main text. Figure S1 contains the lag distributions of other strains from their dataset which highlights a bimodality in lag durations. For Fig. 5b in the main text, we took single-cell lag data for wild yeast strains and evolved strains from Dataset S1 and Dataset S2 of New et al. 2014 (Fig. S2a,b), respectively, and for varying pre-shift durations in glucose from Cerulus et al. 2018 (Fig. S2c), to estimate the lag and fraction of recoverers in a population. Raw single-cell data from [15] presented in Fig. S2c were shared through personal communication by Bram Cerulus and Kevin Verstrepen.

**Fig. S1.**
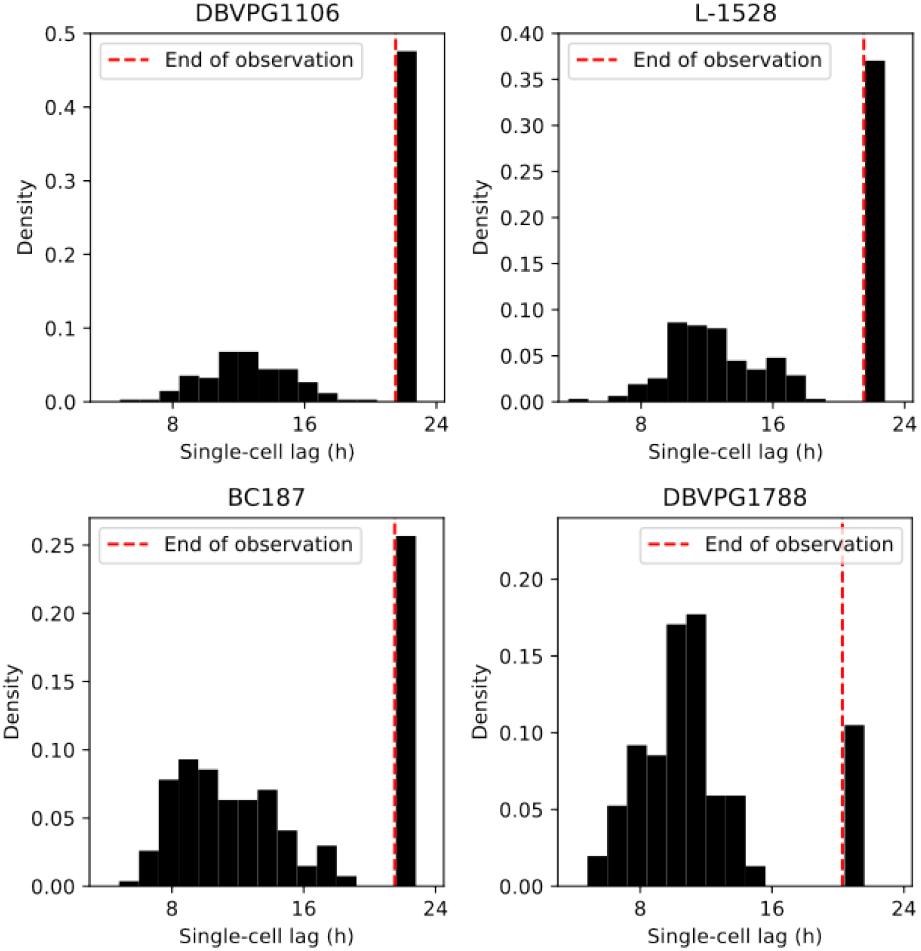
Single-cell lag distributions of different yeast strains.

Most single-cell lag distributions are bimodal, i.e., composed of a group of recoverer cells that recover from lag and arrester cells which do not resume growth until the end of the experiment. We compute the mean lag duration of the recoverer population for each strain and plot it against their fraction in the population.

## Appendix B: Cellular decision-making framework

### 1. Phenotype heterogeneity in fluctuating environments

We consider a scenario where a cellular population experiences transitions between a preferred environment (P) and a non-preferred environment (NP). Cells can take any one of *K* possible phenotypes. For example, a cell with a phenotype that allows for rapid growth in P may experience a longer lag phase or greater death rate when the environment switches to NP. In P, cells grow and rapidly switch their phenotypes so as to maintain a phenotype distribution, ***ϕ*** = (*ϕ*_1_, *ϕ*_2_, …, *ϕ*_*K*_) (with ∑_*k*_ *ϕ*_*k*_ = 1), within the population. The selection of a phenotype distribution could occur through stochastic switching between phenotype states; we discuss this further in Section F. The distinction between P and NP is that cells *cannot* choose their phenotype distribution in NP. This corresponds to the case where cells experience a non-growing phase (lag, starvation, or death) in NP. In this non-growing phase, the population cannot readily express genes and synthesize new proteins, and thus cannot arbitrarily choose its phenotype distribution.

**Fig. S2.**
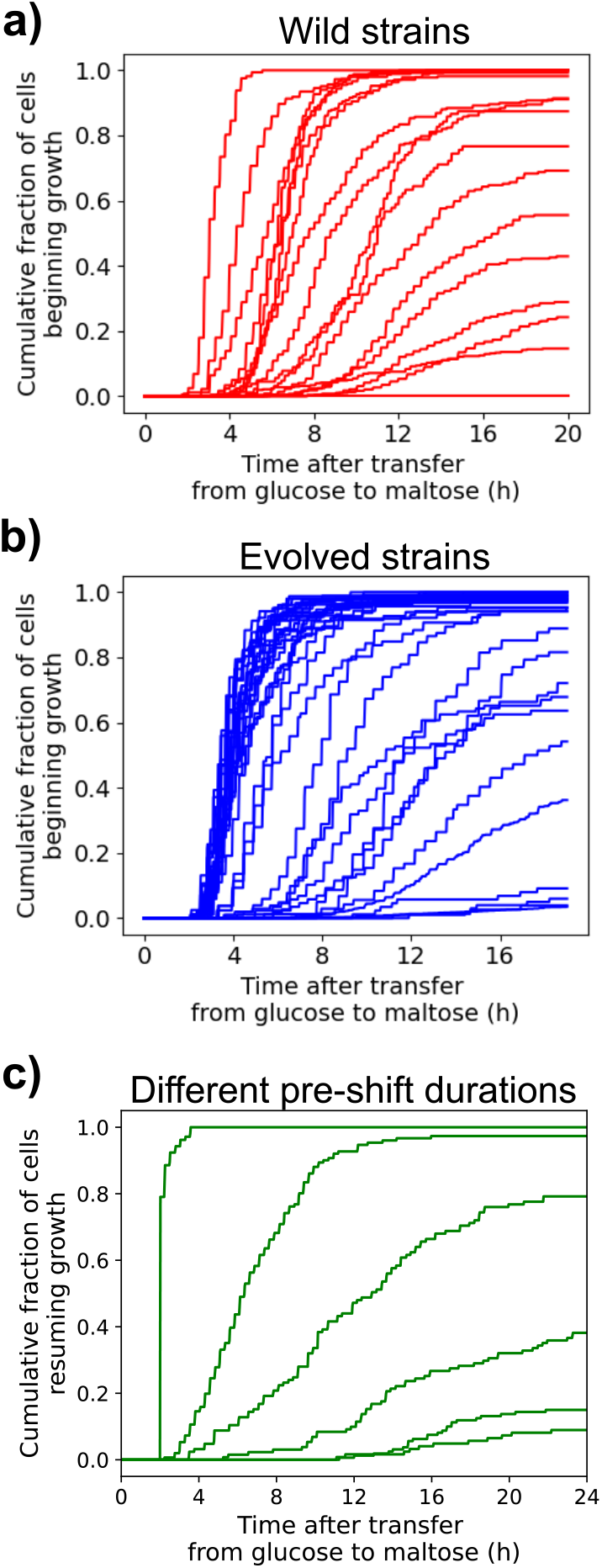
(a) Cumulative fraction of cells resuming growth after the nutrient switch for wild strains of yeast. (b) Cumulative fraction of cells resuming growth after the nutrient switch for evolved populations of yeast. (c) Cumulative fraction of cells resuming growth after the nutrient switch for populations experiencing varying pre-shift durations in glucose.

The state of the environment is a stochastic process. The instantaneous rate at which the population switches from P to NP (denoted *ω*) or from NP to P (denoted *ω*′) could have arbitrary dependence on the history of environmental states (for example, the time since the last switch from NP to P). Cells in P can *anticipate* the transition from P to NP and choose their phenotype distribution ***ϕ*** so as to maximize the long-term growth rate of the population. In the main text, we present a simplified version where cells make decisions that are independent of history. In this simplified version, the environment switches from P to NP and NP to P at constant rates *ω* and *ω*′ respectively. Here, we present a general version of the framework where cellular decisions are allowed to be history-dependent.

### 2. Model setup

We developed a general optimization framework based on dynamic programming to investigate how phenotype heterogeneity influences population fitness under the scenario described in the previous section. This framework allows us to model cellular decision-making using methods similar to those used in reinforcement learning and control theory.

Let *s*(*t*) represent the state at time *t* and *γ*(*s*(*t*), *k*) denote the instantaneous growth rate of a cell with phenotype *k* in state *s*(*t*). *s*(*t*) includes information about whether the current environment is P or NP, and possibly additional information about the growth medium in P (which will affect instantaneous growth rate), the time since the most recent shift from P to NP (which is important to incorporate lag), or historical information the cell uses to better anticipate future environmental transitions. The probability of transitioning from state *s* to *s*′ in interval *dt* is given by the transition probability *P* (*s*(*t* + *dt*) = *s*′|*s*(*t*) = *s*). This transition probability is time-translation invariant, i.e., there is no dependence on absolute time except through *s*. Note that the framework applies for a general state space *s*(*t*), and it can thus capture potentially complex historical dependencies.

Given *s*(*t*) and the phenotype distribution ***ϕ***(*t*), the instantaneous growth rate of a population of *N* cells is

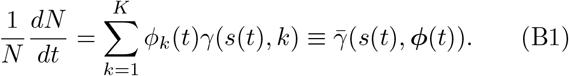

If the initial number of cells is *N*_0_ and the number of cells after time *T* is *N*_*T*_, the long-term growth rate is

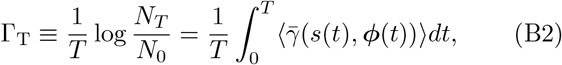

where the angled brackets denote an expectation over changes in environmental state. Our goal is to find the optimal strategy to select ***ϕ*** in P such that Γ_*T*_ is maximized as *T* → ∞.

Instead of directly maximizing 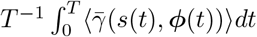 as *T* → ∞, we find the strategy that maximizes 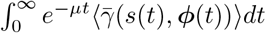 as *µ* → 0. This alternative objective is commonly used to remove time as a state variable when computing the optimal strategy. No singularities arise in the limit *µ* → 0 as the decision to pick a particular ***ϕ*** matters only until the next time the population encounters P.

A population in P selects its phenotype distribution ***ϕ*** at every interval *dt* (where 1*/dt* is much larger than the typical growth rate of the cells in P). We now derive our central optimization objective. We write down a (Bell-man) dynamic programming equation for the long-term growth rate *given* that the population is in state *s* in environment P, assuming the population makes the best possible choice for the phenotype distribution ***ϕ***. Let’s call this optimal long-term growth rate Γ_P_(*s*). We have

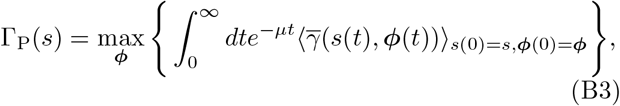

where the expectation (represented by the angled brackets) is over possible future states given the current state *s*(0) = *s* and the phenotype distribution ***ϕ***. Splitting the integral,

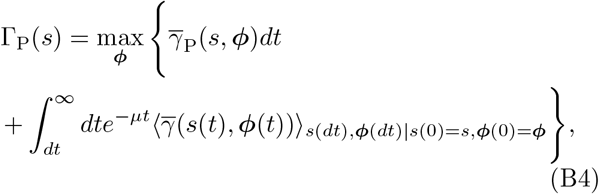

where the expectation is now over the state of the population at *dt* given that at *t* = 0 it is at *s*(0) = *s*, ***ϕ***(0) = ***ϕ***. Setting *t* = *t*′ + *dt*, we have

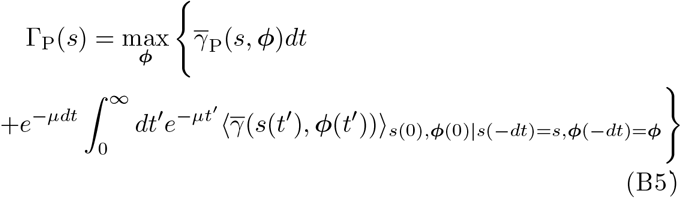

where we used the fact that the transition matrix *P* (*s*(*t*+ *dt*), ***ϕ***(*t* + *dt*)|*s*(*t*), ***ϕ***(*t*)) is time-translation invariant.

Given that the population is in P at state *s*, there are two possibilities: after *dt*, either the environment stays in P with probability 1 − *ω*(*s*)*dt*, or switches to NP with probability *ω*(*s*)*dt*. If the environment stays in P and the state transitions to *s*′, then the expected long-term growth rate after *dt* is Γ_P_(*s*′). If the environment transitions to NP and the state transitions to *s*′, we will denote the expected long-term growth rate by Γ_NP_(*s*′, ***ϕ***) (which we specify further below). We split the integral in the above equation based on the probability of these two possibilities,

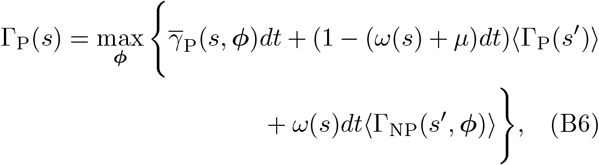

where we have dropped the subscripts specifying that the expectations are over the state and phenotype distribution at the subsequent time step, i.e., *s*′, ***ϕ***′, given *s* at the current time step. Importantly, the choice of ***ϕ*** does not matter if the environment stays in P. Moreover, the term involving ⟨Γ_P_(*s*′)⟩ can be ignored when computing the optimal strategy. To obtain the optimal strategy ***ϕ***\*, we simply have

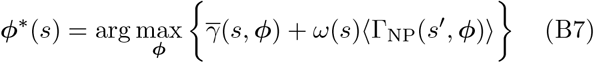

Now, we unpack Γ_NP_(*s*′, ***ϕ***). Let Π(*τ*) be the probability density that the environment stays in NP for duration *τ* and let *s*^′′^ be the state that the environment ends up in after the switch back to P after interval *τ*. We have

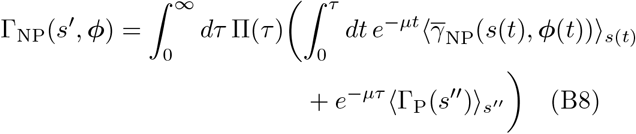

where *s*(*t*) and ***ϕ***(*t*) are now the state and distribution of phenotypes at time *t* after the switch to NP given that the state and distribution were *s*′ and ***ϕ*** when the switch occurred. The first term on the right hand side does depend on *µ*, but it does not contribute to the calculation of ***ϕ***\* if Π(*τ*) decays to zero on timescales much smaller than 1*/µ*. The second term on the right hand side contains ⟨Γ_P_(*s*^′′^)⟩ which is the expected long-term growth rate once the environment switches back to P. This term does not depend on ***ϕ*** and can therefore be ignored when finding the optimal ***ϕ***. We arrive at

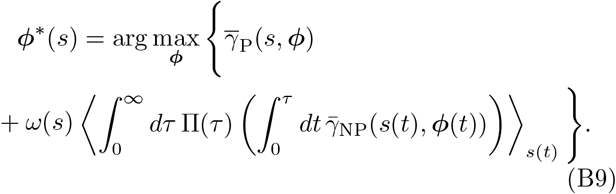

Interchanging integrals and defining 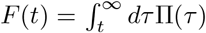, we re-write the above as

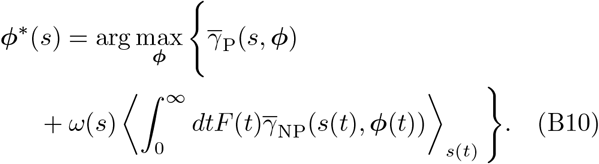

The average growth rate after time *t* in NP is

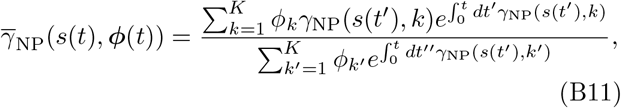

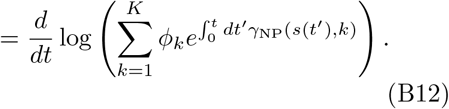

Plugging this expression into Eq. B10 and integrating by parts, we finally obtain the objective

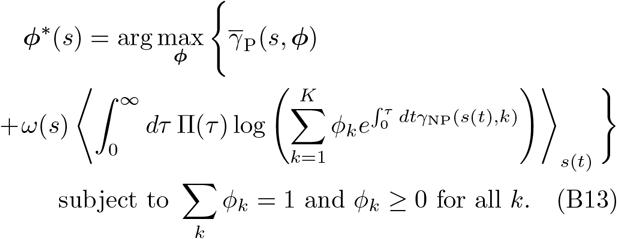

This expression has an intuitive interpretation: the optimal strategy balances the instantaneous growth rate in P and the expected long-term growth rate if the environment were to transition to NP. We now show how this framework can be used to find the optimal decision-making strategy in two scenarios involving a growth-lag trade-off and a growth-death trade-off.

### 3. Balancing growth and lag

Equation B13 is applicable generally when cells transition between preferred and non-preferred environments. In this section, we discuss the situation where a population of cells experiences rapid growth in the preferred environment (P) and undergoes lag phase before resuming growth in the non-preferred environment (NP). In P, cells grow such that *γ*(P, *k*) = *γ*_*k*_. In NP, cells experience lag before they can resume growth: *γ*(*τ*_NP_, *k*) = *γ*′Θ(*τ*_NP_ − *ℓ*_*k*_), where *τ*_NP_ is the time spent in NP since the last switch from P and *ℓ*_*k*_ is the lag of phenotype *k*. We consider 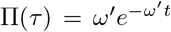 and also ignore any other dependence on the state *s*. Interchanging the integrals over *τ* and *t* in Eq. B13, we get

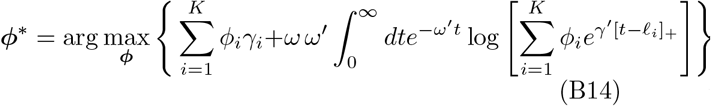

Making substitutions: 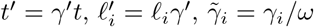, and *η* = *γ*′*/ω*′, we now have

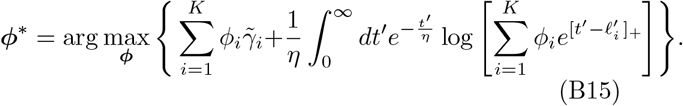

### 4. Balancing growth and death

In an alternative paradigm, rather than experiencing a lag phase in the non-preferred environment (NP), cells are subjected to lethal conditions and die at phenotype-specific rates. In this framework, phenotype variation is characterized not by differences in lag durations, but by distinct death rates *ξ*_*k*_ associated with each phenotype in NP. In P, cells grow such that *γ*(P, *k*) = *γ*_*k*_ and in NP, cells die such that *γ*(NP, *k*) = *ξ*_*k*_, where *ξ*_*k*_(≤ 0) is the death rate for phenotype *k*. Similar to the previous section, we use Eq. B13 and modify the second term concerned with long-term growth in NP,

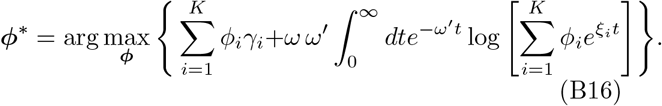

After making substitutions: 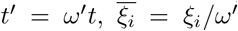 and 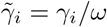, we have

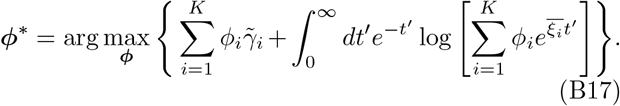

We determine the optimal phenotype distribution ***ϕ***\* numerically.

### 5. Numerical optimization

We assume the population contains a distribution over *K* phenotypes, where phenotype *k* (where *k* takes on values from 1 to *K*) has growth rate *γ*_*k*_ (*γ*_min_ ≤ *γ*_*k*_ ≤ *γ*_max_) in P. In the growth-lag scenario, phenotypes are characterized with lags *ℓ*_*k*_ in NP and all phenotypes have the same growth rate *γ*′ in NP when cells recover from lag phase. In the growth-death scenario, phenotypes are characterized with death rates *ξ*_*k*_ in NP. The phenotype distribution is given by ***ϕ*** = (*ϕ*_1_, .., *ϕ*_*k*_, .., *ϕ*_*K*_).

Step-by-step procedure to run the optimization scheme:

- First we pick the parametric form for the trade-off between growth and lag (or death). This gives us growth rates {*γ*_*i*_} in P and the corresponding lag durations {*ℓ*_*i*_} (or death rates {*ξ*_*i*_}) in NP.
- Choose transition rates *ω* and *ω*′.
- Initialize ***ϕ*** = {*ϕ*_*i*_} such that *ϕ*_*i*_ = 1*/K*. To prevent having to enforce an extra constraint of 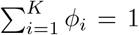 during optimization, we parametrize the probabilities as 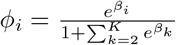 for 2 ≤ *i* ≤ *K* and 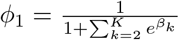. Here {*β*_*i*_} are unconstrained parameters that we can directly optimize over.
- Using the Adam optimizer in the Optax library applied to a JAX implementation in Python, we numerically find {*β*_*i*_} that maximizes Γ_P_(***ϕ***) from Eq. B15. We then obtain the optimal phenotype distribution ***ϕ***\*.

## Appendix C: Optimal phenotype distribution consists of a few discrete phenotypes

We consider our optimization problem where the population selects a phenotype distribution ***ϕ*** to maximize a long-term growth objective of the general form

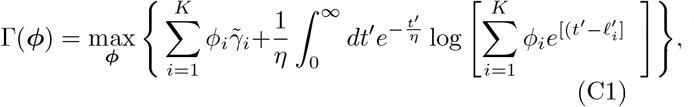

with the constraints:

- *ϕ*_*i*_ ≥ 0 ∀*i*
- 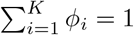

For this constrained nonlinear optimization problem, we use the Karush-Kuhn-Tucker (KKT) conditions to make predictions about the structure of the optimal solution. We construct the corresponding Lagrangian function,

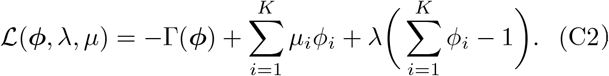

for some Lagrange multiplier *λ* and KKT multipliers *µ*_*i*_ ≥ 0. To maximize Γ(***ϕ***) given the constraints, the stationarity condition requires

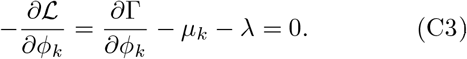

Let 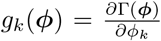 denote the marginal value of phenotype *k*. Then the support of the optimal solution (i.e., the set of phenotypes with *ϕ*_*k*_ *>* 0) is determined by the set of indices where *g*_*k*_ = *λ*. We get this from the complementary slackness condition which requires that *µ*_*i*_*ϕ*_*i*_ = 0 for *i* = 1, 2, …, *K*. Geometrically, this corresponds to the intersection points of a horizontal line at height *λ* with the graph of *g*_*k*_ as a function of phenotype lag 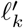. Let’s examine the explicit form of *g*_*k*_:

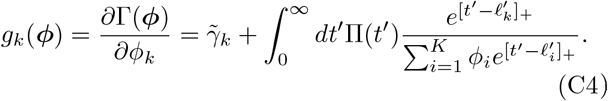

In our framework, the rescaled lag durations *ℓ*′ and preferred-environment growth rates 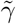 are strictly ordered such that: 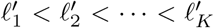 and 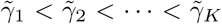. This implies that the first term in *g*_*k*_(***ϕ***) monotonically increases with *k*. In the second term, the dwell time distribution Π(*t*′) acts as a discount factor such that for larger *t*′, Π(*t*′) is smaller since the distribution monotonically decreases with *t*′. This implies that the contribution to the integral for larger 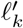 is reduced which in turn means that the second term in *g*_*k*_(***ϕ***) monotonically decreases with *k. g*_*k*_(***ϕ***) is composed of the sum o two terms with opposing monotonicities. If *g*_*k*_ is strictly monotonic in 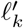, then the horizontal line intersects the curve at most once, implying that the optimal strategy is supported on a single phenotype. The number of times *g*_*k*_ intersects with the horizontal line determines the number of phenotypes with non-zero probability. For well behaved Π(*t*), we expect at most a few discrete pheno types, though the precise number depends on Π(*t*) and the growth-lag trade-off.

### Single timescale in NP

In this section, we present a geometric argument for why we expect at most two phenotypes when the distribution of dwell times in NP is unimodal. This geometric (C1) argument complements the formal argument in the main text using the KKT theorem. Suppose that the time spent in NP is a constant *t* instead of being exponentially distributed. We have for *K* phenotypes (with arbitrary growth rates)

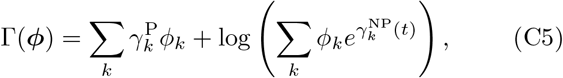

where 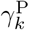 is the growth rate in P and 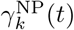 is the effective growth rate in NP for phenotype *k*. In the growth-lag scenario 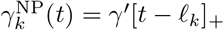, while in the growth-death scenario 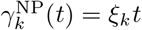.

Consider ***ϕ*** such that 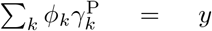 and 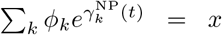. The two constraints *ϕ*_*k*_ ≥ 0 and ∑_*k*_ *ϕ*_*k*_ = 1 define a closed feasible region in ***ϕ***. This closed feasible region in ***ϕ*** translates to a closed feasible region *D* in the two-dimensional (*x, y*) plane. Consider the family of curves *y* = Γ − log *x* parameterized by Γ. As Γ is decreased from infinity, the optimal feasible solution is due to the curve that just touches *D*. Now we show that the boundary points of *D* correspond to *ϕ* such that at most two of the phenotypes have non-zero probability mass.

First note that the intersection of the hyperplane 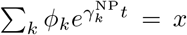 with the probability simplex *ϕ*_*k*_ ≥ 0, ∑_*k*_ *ϕ*_*k*_ = 1 is a convex hull formed by points on line segments joining two vertices of the probability simplex. These are points that have non-zero probability for at most two of the phenotypes. Next, for a given value of *x*, the top and bottom boundaries of *D* correspond to the maximum and minimum value of *y* taken over this convex hull. However, the maximum and minimum of *y* is always achieved at the vertices of the convex hull. The optimal solution therefore is such that at most two of the phenotypes have non-zero mass.

While this argument does not generalize to arbitrary distributions over *t*, it helps rationalize why at most two phenotypes are largely sufficient when the distribution over *t* has one typical timescale (such as for an exponential distribution).

## Appendix D: Under what scenarios is a bimodal solution favored?

### 1. Growth–lag paradigm

We aim to obtain a condition on the curvature of the growth-lag trade-off relation *γ*(*ℓ*) for when the one-state solution splits into a two-state solution. We first obtain the best one-state solution and then ask if there is a better two-state solution such that the two states have lags that are in an *ε*-neighborhood of the best single-state solution.

Consider the general two-state objective function

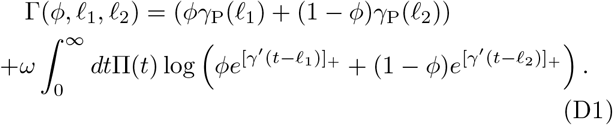

We set *ω* = *γ*′ = 1 noting that growth rates in P are measured in units of *ω* and time spent in NP and lag durations are measured in units of 1*/γ*′. The optimal single-state solution (with *ϕ* = 1 in Eq. D1 denoted by Γ_1_ is obtained by maximizing

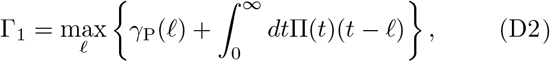

which leads to the optimality condition and single-state growth rate

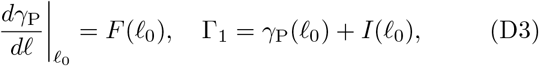

where *ℓ*_0_ is the lag that maximizes the rhs of Eq. D2, and we denote 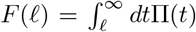 and 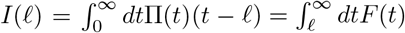 (the latter is obtained via integration by parts). We only consider cases where the best single-state lag *ℓ*_0_ is finite, that is, Γ_1_ *> γ*_P_(∞).

We now calculate the best two-state solution such that *ℓ* −*ε* ≤ *ℓ*_1_, *ℓ*_2_ ≤ *ℓ*_0_ +*ε* where *ε* is a small positive constant. To do this, we expand each term in Eq. D1 to second order. Define *ε*_1_ = *ℓ*_1_ − *ℓ*_0_, *ε*_2_ = *ℓ*_2_ − *ℓ*_0_. All *γ* terms and Taking the maximum over *ϕ* in Eq. D9, we get for the its derivatives are evaluated at *ℓ*_0_. The first term on the

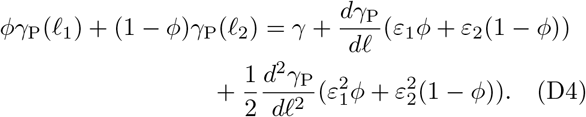

We split the second term to three parts

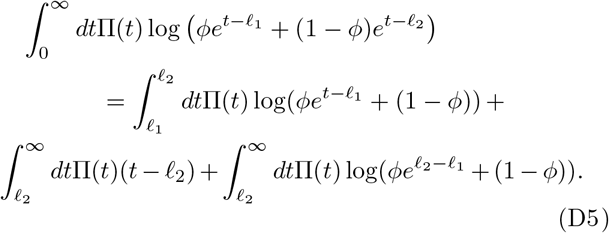

The first term in Eq. D5 to second order in *ε* is

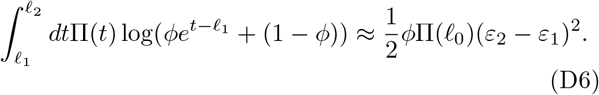

The second term in Eq. D5 to second order in *ε* is

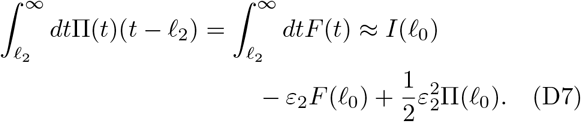

The third term in Eq. D5 to second order in *ε* is

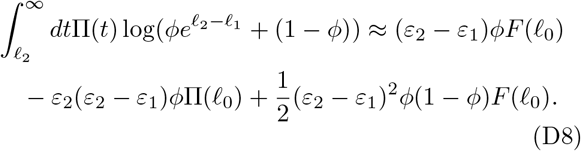

Collecting the zeroth order terms, we get *γ*_P_(*ℓ*_0_) + *I*(*ℓ*_0_), which is precisely Γ_1_. The first order terms cancel out exactly once we use the stationarity condition *dγ*_P_*/dℓ* = *F* (*ℓ*_0_) at *ℓ*_0_. After adding up the contributions from the second order terms, the two-state objective Γ_2_ to second order in *ε* is

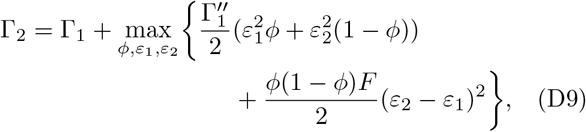

where 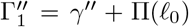 is the second derivative of the single-state objective at *ℓ* = *ℓ*_0_, *F* = *F* (*ℓ*_0_) and the optimization has constraints 0 *< ϕ <* 1 and |*ε*_1_|, |*ε*_2_| ≤ *ε*. Moreover, *ε*_1_ ≠ *ε*_2_ (since otherwise that would be a single-state solution).

Taking the maximum over *ϕ* in Eq. D9, we get for the optimum *ϕ*\*,

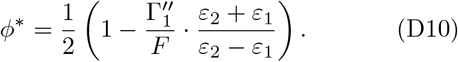

Note that 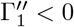 since *ℓ*_0_ is an interior point that attains the maximum of the single-state objective. Denote 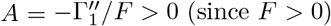. Plugging *ϕ*\* into Eq. D9 and simplifying, we get

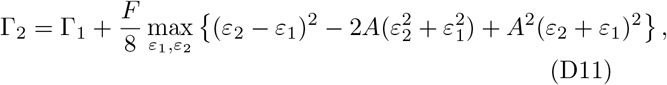

with the constraints *ε*_1_ ≠ *ε*_2_ and |*ε*_1_|, |*ε*_2_| ≤ *ε*. There are three possibilities for the optimal *ε*_1_, *ε*_2_: (1) they are both interior points, i.e., they belong to the open interval (−*ε, ε*), (2) one of them takes a boundary value of ±*ε* and the other is an interior point, or (3) both of them are at the boundary, in which case without loss of generality *(ε*_2_ = −*ε*_1_ = *ε)* (since they cannot be equal). We now consider each case separately.

If they are both interior points, we can take the derivatives of Eq. D11 w.r.t *ε*_1_ and *ε*_2_ and equate them to zero. We get

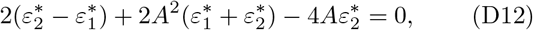

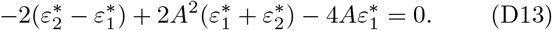

Solving for the two we get 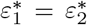, which violates the *ε*_1_ ≠ *ε*_2_ constraint.

The second case is that one of them is on the boundary say 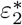. Then, taking the derivative of Eq. D11 w.r.t *ε*_1_ and setting it to zero, we get

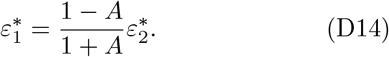

The term being maximized in (34) is then

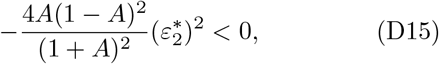

since *A >* 0. The single-state solution outcompetes the best two-state solution in this scenario.

This leaves the third case that *ε*_2_ = −*ε*_1_ = *ε*, which gives for the term being maximized in Eq. D11 as 4(1− *A*)*ε*^2^. Note that *ϕ*\* = 1*/*2 in this case which satisfies the constraint 0 ≤ *ϕ* ≤ 1. When *A <* 1, the two-state solution outcompetes the single-state solution. Thus,

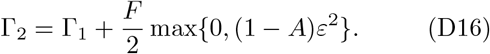

The transition from a single-state to two-state solution occurs at *A* = 1, or in terms of the trade-off and switching statistics,

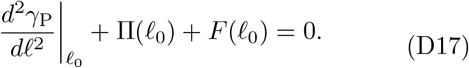

Note that this is a sufficient condition, but not a necessary one.

#### a. Test for critical curvature

For a chosen growthlag trade-off with form 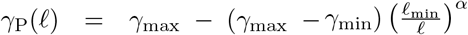, we identify the single-phenotype optimum (*ℓ*_0_, *γ*_0_). At this point, we construct small parabolas with varying curvatures to test when the solution transitions from a single-state solution to a two-state solution. After performing a rotation, the parabola with vertex (*ℓ*_0_, *γ*_0_) is given by

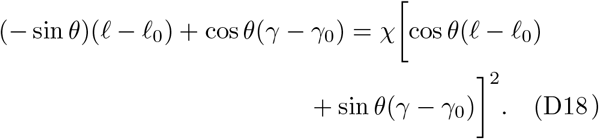

Taylor expanding *γ* around *γ*_0_ and substituting in the equation above, we get

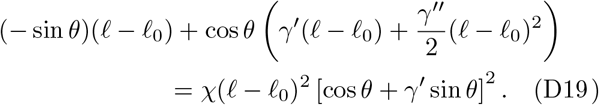

Since *γ*′ = sin *θ/* cos *θ*, this equation reduces to

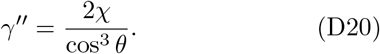

In the previous section, we derived the condition for the transition from a single-state solution to a two-state solution, − [*γ*^′′^ + Π(*ℓ*_0_)]*/F* = 1. Assuming Π(*t*) = *e*^−*t*^, the condition becomes

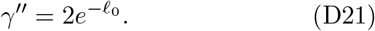

Finally, by equating Eq. D20 and Eq. D21, and noting that 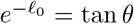, we get a relation for critical curvature parameterized by *χ*_crit_ for the transition

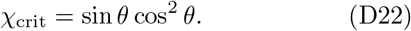

We confirmed this result numerically which is described in Fig. S3a,b.

#### b. Power-law growth-lag trade-off and exponential dwell times in NP

When *γ*(*ℓ*) = −*ℓ*^−*α*^ and 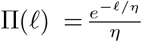 (recall *η* = *γ*′*/ω*′), *ℓ*_0_ is obtained by solving

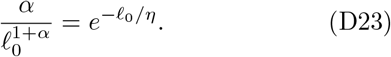

Using Eq. D17, we have

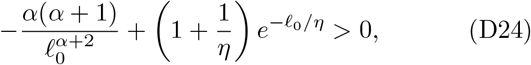

which gives

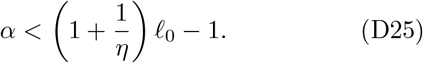

Plugging back *η* = *γ*′*/ω*′ and reintroducing units, *ℓ*_0_ → γ′*ℓ*_0_, we have

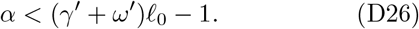

### 2. Growth–death paradigm

We now consider a similar analysis for the growth-death paradigm. We analyze two cases that are distinguished by the sign of the curvature of the growth-death trade-off. In the first case, we show that the optimal phenotype distribution *always* consists of at most two specialist phenotypes, regardless of the distribution of dwell-times in NP. In the second case, analogous to our analysis in the growth-lag scenario, we obtain a sufficient condition on the curvature of the growth-death trade-off that favor a heterogeneous, specialist strategy compared to a generalist strategy.

Let *R* denote the set of physiologically plausible *γ, ξ* values (recall that the *γ* ≥ 0, *ξ* ≤ 0). We define the boundary ∂*R* of this region as those phenotypes for which there is no other phenotype in *R* where both *γ* and *ξ* are larger.

**Fig. S3.**
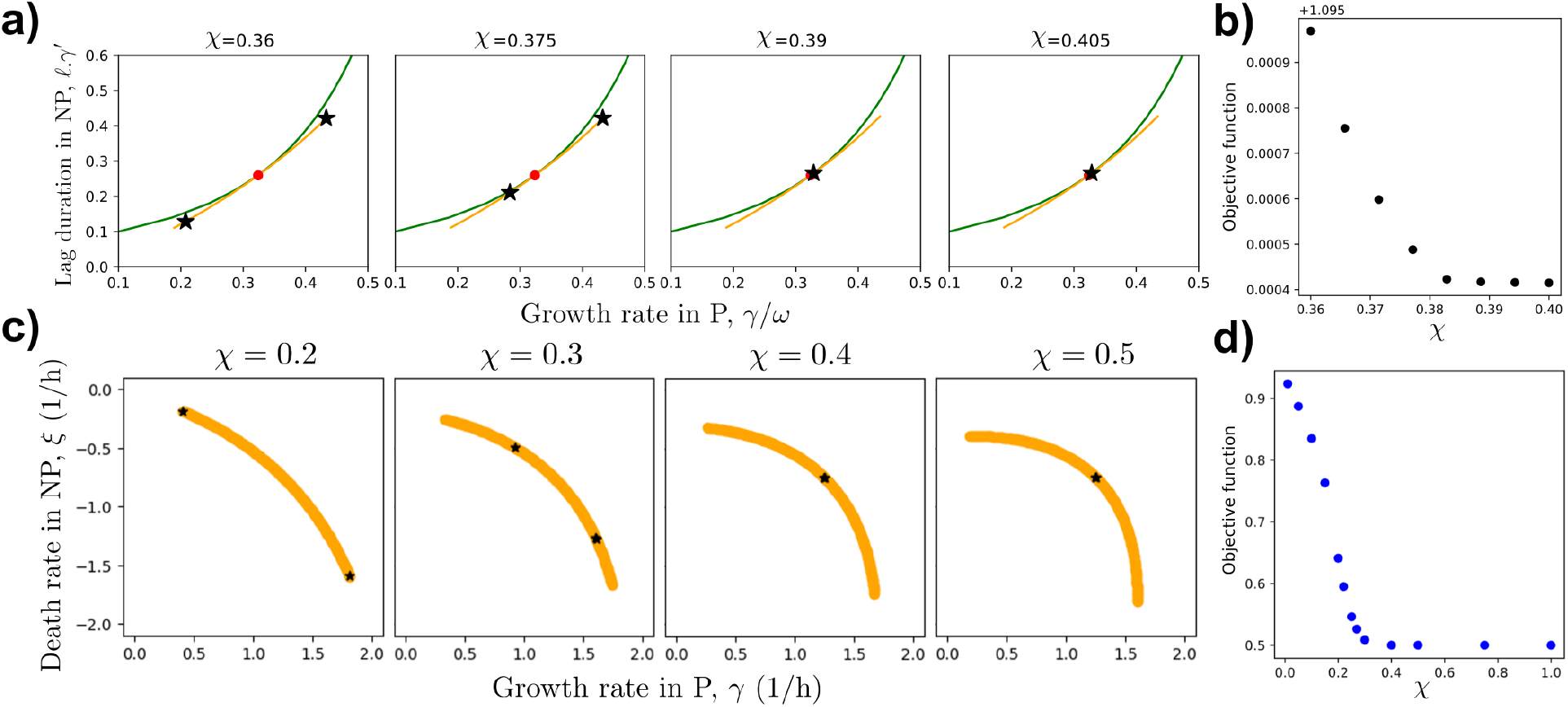
Transition from a two-state solution to a single-state solution in the growth-lag system (a) and (b), and growth-death system (c) and (d). (a) The green curve represents the original power-law growth-lag trade-off for which the single-phenotype optimum, (*γ*_0_, *ℓ*_0_), is represented by the red circle. Taking this point as the vertex, we construct small parabolas with varying curvatures parameterized by *χ* and numerically determine the optimal solution. For a specific value of *χ* = *χ*_crit_, there is a transition from a two-state solution to a single-state solution. For the example shown above, this transition occurs close to the analytic prediction of *χ*_crit_ = 0.3822. (b) Maximum long-term population growth rate estimate for parabolic trade-offs with different curvatures, *χ* in growth-lag system. (c) Analogous to (a): Optimal solutions for different small parabola growth-death trade-offs with varying *χ* are tested. The analytic prediction for *χ*_crit_ is 0.3535 for this example. (d) Maximum long-term population growth rate estimate for parabolic trade-offs with different curvatures, *χ* in growth-death system.

#### Case 1

We first consider *R* such that there are no physiologically plausible phenotypes on the line joining any two phenotypes on ∂*R*. In this case, we can show that the optimal solution is at most bimodal, regardless of the distribution over intervals in the NP environment.

Consider *K* phenotypes on ∂*R* such that *γ*_1_ *> γ*_2_ *>* · · · *> γ*_*K*_ and *ξ*_1_ *< ξ*_2_ *<* · · · *< ξ*_*K*_. We have

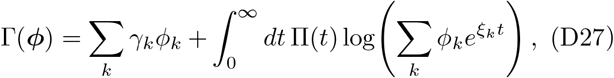

where Π(*t*) is the distribution over intervals in NP. Say ∑_*k*_ *γ*_*k*_*ϕ*_*k*_ = *x*. We will show that for each *x* that is feasible, the optimal solution is a mixture of the 1st and *K*th phenotypes.

The feasible points given ∑_*k*_ *γ*_*k*_*ϕ*_*k*_ = *x* is the convex hull formed by points on the edges of the probability simplex. Let the vertices of the convex hull be ***q***_1_, ***q***_2_, …, ***q***_*L*_. Any feasible point *ϕ* (given *x*) can be written as ∑_*µ*_ *u*_*µ*_***q***_*µ*_ where *u*_*µ*_ ≥ 0, ∑_*µ*_ *u*_*µ*_ = 1. Consider the vector ***w***(*t*) whose *k*th component is 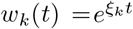. The last term in Eq. D27 can then be written as 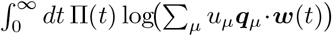.

Whenever the hyperplane ∑_*k*_ *γ*_*k*_*ϕ*_*k*_ = *x* intersects the probability simplex (*γ*_1_ ≥ *x* ≥ *γ*_*K*_), it always intersects the edge joining the vertex *ϕ*_1_ = 1 and the vertex *ϕ*_*k*_ = 1 Say this point of intersection is ***q***_1_. We have

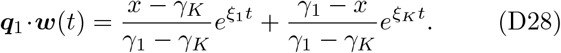

We will show that ***q***_1_ · ***w***(*t*) ≥ ***q***_*µ*_ · ***w***(*t*) for all *µ* and *t*. This implies that for every feasible *x*

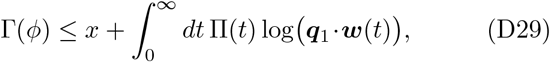

with equality when *u*_1_ = 1, i.e. when all probability mass is on the 1st and *K*th phenotypes.

To prove ***q***_1_ · ***w***(*t*) ≥ ***q***_*µ*_ · ***w***(*t*), first consider an intermediate phenotype *i* with growth rates *γ*_*i*_, *ξ*_*i*_. Let ***q***_*µ*_ be the intersection of the hyperplane with the line joining *ϕ*_1_ = 1 and *ϕ*_*i*_ = 1 (*γ*_1_ ≥ *x* ≥ *γ*_*i*_):

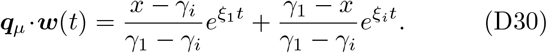

A short calculation gives

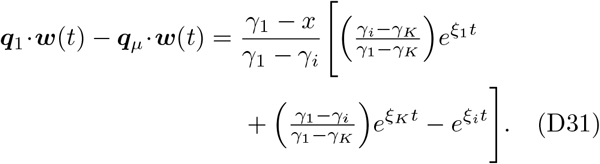

Since *e*^*t*^ is convex, one obtains

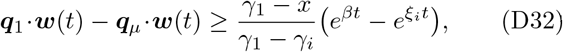

where

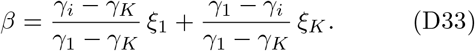

Thus ***q***_1_ ·***w***(*t*) ≥ ***q***_*µ*_ ·***w***(*t*) if

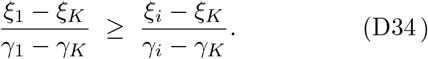

By the assumed shape of ∂*R*, this holds for all *i*. A similar argument applies when the hyperplane intersects the edge joining *ϕ*_*k*_ = 1 and *ϕ*_*i*_ = 1 (*γ*_*K*_ ≤ *x* ≤ *γ*_*i*_). Hence ***q***_1_ ·***w***(*t*) ≥ ***q***_*µ*_ ·***w***(*t*) for every *µ*.

#### Case 2

Now suppose ∂*R* does not satisfy the conditions of Case 1, so solutions with more than two modes become possible. Nonetheless, we can state when more than one phenotype is favored. The best single-phenotype strategy corresponds to the point where a line of slope −1 is tangent to ∂*R* in the (*γ, ξ*) plane. If the curvature of ∂*R* near this point is small enough, a heterogeneous strategy outcompetes the single-phenotype strategy.

To see this, consider an infinitesimal parabola *w* = *χz* around the best phenotype (*γ, ξ*). The *z*-axis is along the tangent; the *w*-axis points away and downward. We look at solutions on this parabola with |*z*| ≤ *ε* ≪ 1 for various *χ*.

As in Case 1,

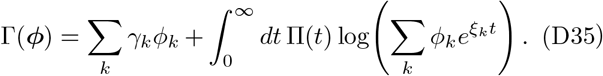

Rescale *t* and *ξ*_*k*_ so that ⟨*t*⟩ = 1. Since solutions are at most bimodal, let one phenotype have probability *ϕ* and lie at *z*_1_, the other at *z*_2_. Writing *γ*_*i*_ = *γ* + Δ*γ*_*i*_ and *ξ*_*i*_ = *ξ* +Δ*ξ*_*i*_ for *i* = 1, 2, a second-order Taylor expansion gives

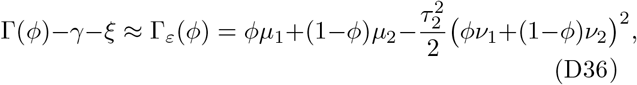

with 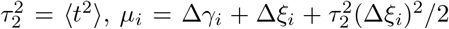 and *ν*_*i*_ = Δ*ξ*_*i*_. Optimizing over *ϕ* yields

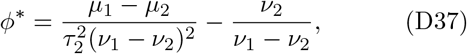

for 0 ≤ *ϕ*\* ≤ 1. Substituting back,

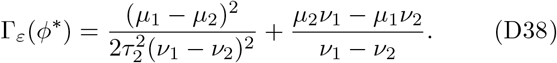

With 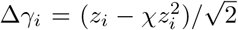 and 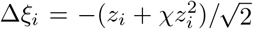 and retaining terms up to second order,

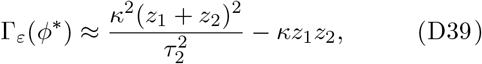

where 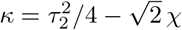.

For *κ <* 0, *a >* 1 and the optimum reduces to the single best phenotype; in this case,

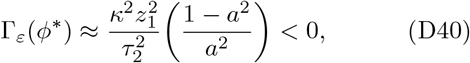

which is suboptimal.

If *κ >* 0, a bimodal solution with *z*_1_*z*_2_ *<* 0 beats the single-phenotype solution (provided 0 ≤ *ϕ*\* ≤ 1). One shows the optimum occurs at *z*_1_ = −*z*_2_ = *ε*:

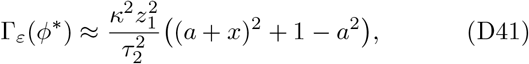

where *x* = *z*_2_*/z*_1_ and 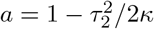. Since *a <* −1 when *κ >* 0, *x* = −1 maximizes Eq. D41. Then *ϕ*\* = 1*/*2 and 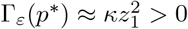, maximized at *z*_1_ = *ε*.

Hence, a heterogeneous solution is favored if *κ >* 0, i.e.,

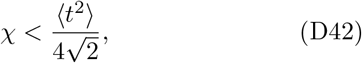

where ⟨*t*^2^⟩ is the second moment of the NP dwell-time distribution. If the curvature *χ* of ∂*R* is below this threshold, a mixed two-phenotype strategy (specialists) outperforms the single-phenotype strategy (generalist). This is confirmed numerically where we plot the maximum long-term growth rate for different *χ* values as shown in Fig. S3d. For *χ* above the threshold, the optimal growth rate stays constant as the single phenotype solution doesn’t change with *χ*. For *χ* below the threshold, we see that higher values for the long-term growth are achievable with two-phenotype strategies.

We chose parabolic curves centered at a specific point with different curvatures *χ* for the growth-death trade-off. In Fig. S3c, as *χ* increases, the optimal solution transitions from a bimodal phenotype distribution to a single-phenotype distribution.

## Appendix E: Two-phenotype system in the growth-lag paradigm

### 1. Why do arresters have infinite lag?

Assuming that the population can have two phenotypes with lags *ℓ*_1_ and *ℓ*_2_ such that *ℓ*_*min*_ ≤ *ℓ*_1_ *< ℓ*_2_ and the fraction of the population with lag *ℓ*_1_ in P is *ϕ*. We consider growth-lag trade-offs such that the growth rate in P of a cell with lag *ℓ* in NP is

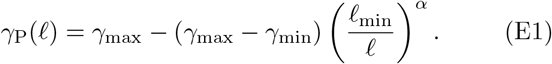

Consider the term to be maximized over *ϕ* in Eq. B15, which we denote as Γ(*ϕ, ℓ*_1_, *ℓ*_2_). We consider Π(*t*) = *ω*′*e*^−*ω*^′^*t*^. Interchanging the integrals over *τ* and *t*, we get

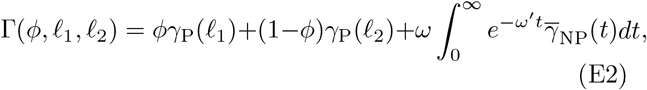

where 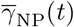 is the average growth rate at time *t* after the environment is switched from P to NP given that the population began with fraction *ϕ* cells in phenotype 1. Specifically,

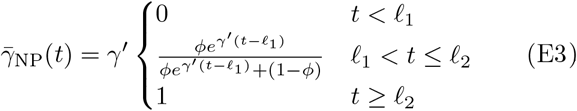

Rescaling time: *t* → *γ*′*t*, and defining *η* = *γ*′*/ω*′, we have

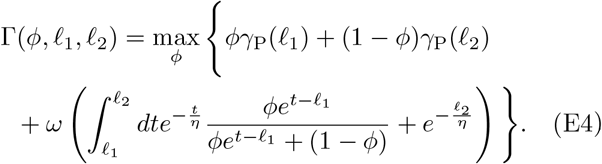

Shifting time as *t* → *t* − *ℓ*_1_ and after a few simplifications, Γ reduces to the objective function

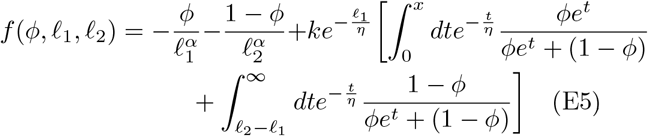

where *k* is a positive constant obtained by absorbing *ω* with 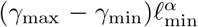, and *ℓ*_2_ = *ℓ*_1_ + *x*. Substituting 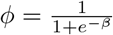, we derive an equivalent form for the objective function

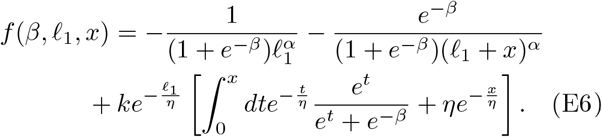

Taking the derivative of *f* with respect to *x* yields

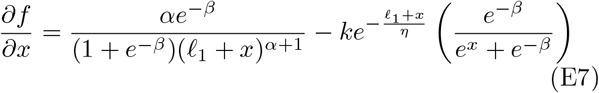

The first term on the right hand side is always positive and decays with *x* as a power-law, whereas the second term is always negative and decays exponentially. For sufficiently large *x*, the first term dominates and the gradient is positive. That is, it is preferable to indefinitely increase *ℓ*_2_. While this argument does not prove that the arrester solution is the preferred phenotype under all scenarios, it helps rationalize why an arrester phenotype with an infinite lag is selected.

### 2. Relating the lag duration of recoverers and the fraction of recoverers

We now consider a situation where the population can choose the fraction of recoverers *ϕ* and their lag duration *ℓ* ≥ *ℓ*_min_ (and consequently their growth rate *γ*(*ℓ*)). This scenario corresponds to *ℓ*_1_ = *ℓ* and *ℓ*_2_ = ∞ from Section E 1. The average growth in 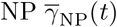 satisfies 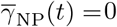 for *t* ≤ *ℓ*. For *t > ℓ*, we have

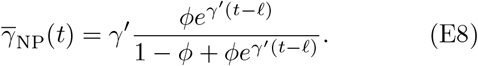

Substituting *t* → *t* − *ℓ* and simplifying, we obtain

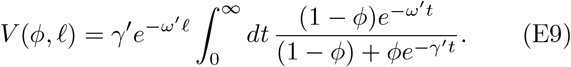

The integral on the right-hand side can be expressed in terms of the hypergeometric function _2_*F*_1_(*a, b*; *c*; *x*). Denoting *η* ≡ *γ*′*/ω*′, we have

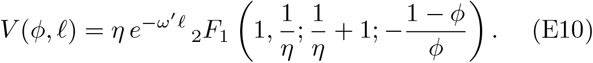

When 0 *< ϕ <* 1 and *ℓ > ℓ*_min_, we can optimize for *ϕ* and *ℓ* by taking the derivative of the growth function and setting it to zero. For a population in state *s* in P, we obtain the optimality relation

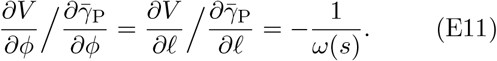

The first equation above expresses a relationship between the optimal *ℓ*\* and *ϕ*\* independent of *ω*(*s*). We then obtain,

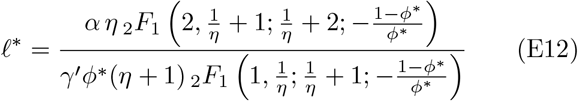

### 3. Inferring the environmental parameters from experimental data

Treating *η* and *α/γ*′ as fitting parameters, we obtained best-fit values of *η* = 1.2 and *α/γ*′ = 21 h (see Fig. 5c). Using the experimental estimate *γ*′ = 0.1 h^−1^ from [16], these values correspond to *α* = 2.1 and *ω*′ = 0.08 h^−1^. To proceed with constructing the phase-portrait for the optimal solution, we use the fact that the maximum growth rate *γ*_P_ on glucose is 0.5 h^−1^. The phase portrait can be seen in Fig. S4. The red region indicates where the yeast system exhibits a bimodal strategy of specialists. This corresponds to *ω* ∈ [0.07, 0.59] h^−1^.

## Appendix F: Switching between two phenotypes in the preferred environment

While setting up the model, we made an assumption that cells rapidly switch between different phenotypes to maintain a constant phenotype distribution in the preferred environment (P). This switching occurs on timescales shorter that any doubling time and typical time spent in P, but slower than the time it takes to switch between P and NP. This ensures that cells cannot adapt while the environment is switching. In this section we illustrate the balance between growth and switching that ensures a constant phenotype distribution in P.

**Fig. S4.**
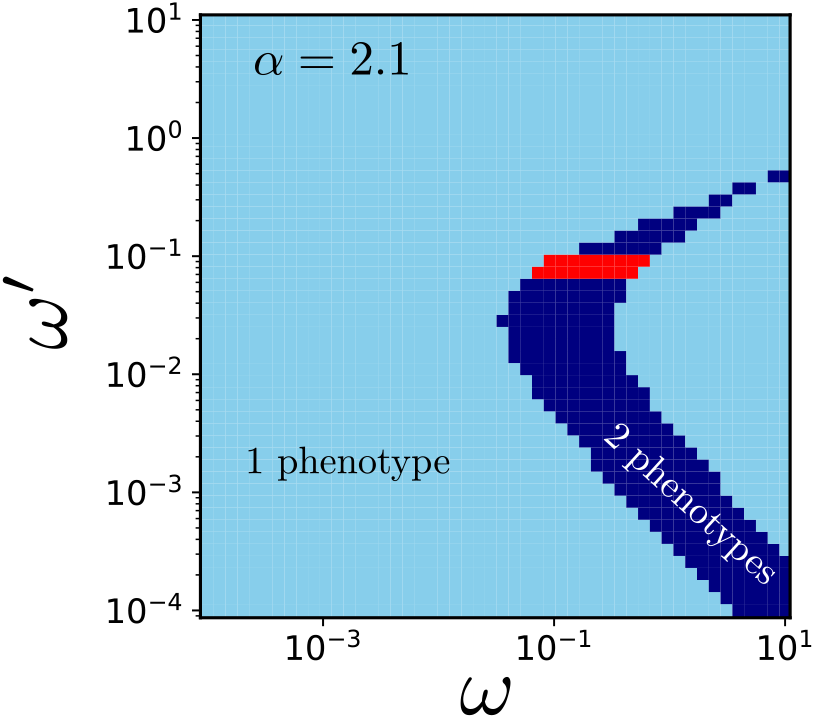
Phase diagram of the optimal number of phenotypes_1_ over the space of environmental switching rates, *ω* and *ω*′, for a trade-off with *α* = 2.1, which is the value inferred from experimental data. The red region corresponds to where the yeast system exhibits a bimodal specialist strategy.

Suppose the population is composed of two phenotypes with growth rates *γ*_1_ and *γ*_2_ (*γ*_2_ *> γ*_1_). *N*_tot_ is the total number of initial cells of which *N*_1_ cells adopt phenotype 1 and *N*_2_(= *N*_tot_ − *N*_1_) cells adopt phenotype 2. The time-evolution of *N*_tot_ and *N*_1_ are governed by

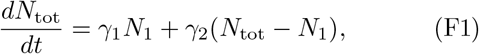

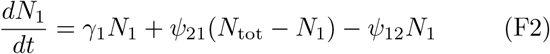

where *ψ*_21_ and *ψ*_12_ are the switching rates from phenotypes 2 to 1 and 1 to 2 respectively. The quantity that remains constant in the preferred environment is *ϕ* = *N*_1_*/N*_tot_ such that *dϕ/dt* = 0. This gives us,

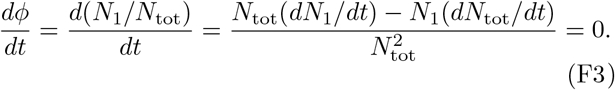

Substituting Eq. F1 and Eq. F2 and after simplifying we get,

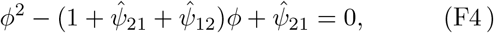

where we have defined 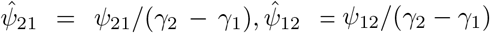. The solution for *ϕ* is given by

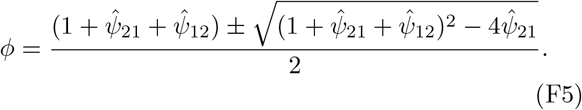

Tuning 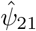 and 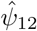 allows for tuning *ϕ* to any value between 0 and 1. For example, when the rate of switching is much larger than the difference in growth rates, i.e., 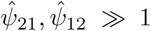, we can approximate the steady-state *ϕ* using Eq. F5 as

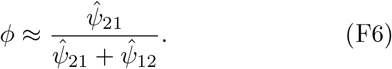

The sum 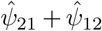 sets the rate at which the population reaches steady-state, whereas the ratio 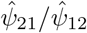 sets the value of *ϕ* at this steady-state.

## Appendix G: A minimal metabolic model illustrating constraints on speed of adaptation

Microorganisms encounter environments in which the available carbon source fluctuates between preferred and non-preferred nutrients. Certain nutrients (e.g. glucose) support rapid growth by activating the corresponding metabolic pathways (e.g. glycolysis), whereas other nutrients (e.g. glycerol, acetate) require induction of alternative pathways (e.g. gluconeogenesis). This results in a characteristic lag phase prior to growth resumption. Fast growth on the preferred nutrient and rapid adaptation (short lag) on the non-preferred nutrient are inherently antagonistic, reflecting a growth–lag trade-off. We present a minimal model, inspired by central metabolism, which captures a growth-lag trade-off that could arise due to antagonism in core pathways.

### Model structure and assumptions

The schematic of the model is shown in Fig. 4a, where:

- *G* and *N* denote the concentrations of the metabolite pools in the preferred environment (P) and non-preferred environment (NP), respectively.
- *g* and *n* denote intracellular concentrations of the central metabolites.
- *r* is the maximal forward flux rate of the irreversible reactions in both P and NP.
- *δ* is the loss rate constant for *g* and *n*.
- *ϕ* and *ζ* are the abundances of the irreversible enzymes that catalyze the forward (*g* → *n*) and reverse (*n* → *g*) reactions, respectively.

We assume Michaelis–Menten kinetics for each enzymatic step. Growth rate linked to biomass production, *µ*, is assumed to depend on the levels of the metabolites *g* and *n*:

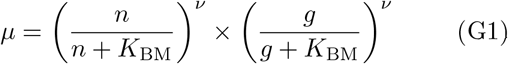

where *K*_BM_ is the half-saturation constant, and the exponent *ν >* 0 captures effective cooperative mechanisms in metabolism.

The time evolution of each metabolite pool is given by mass-action and Michaelis–Menten type terms. In the preferred environment when *G* is present and *N* is absent, the equations are

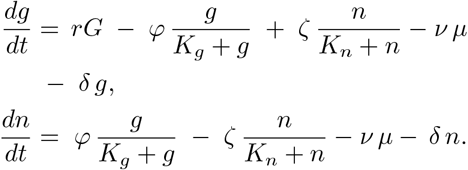

In the non-preferred environment when *G* is absent and *N* is present, the equations are

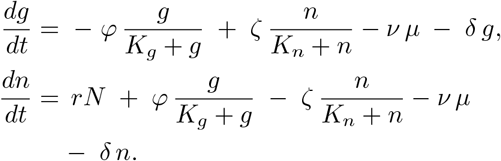

We fix the concentrations of *G* and *N* to be 2 a.u. and 1 a.u. in the preferred and non-preferred environments, respectively.

### Enzyme dynamics and regulation

We model the synthesis and dilution of the irreversible enzymes, *ϕ* and *ζ*, by simple first-order kinetics toward nutrient-specific steady-state levels:

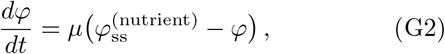

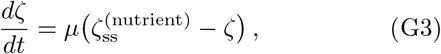

where

- *µ* is the instantaneous growth rate;
- 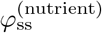 and 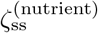 are the steady-state enzyme levels where the ‘nutrient’ may be *G* or *N*;
- 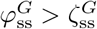 biases flux in the direction *g* → *n* when *G* is the nutrient, and vice versa after a nutrient shift to *N* being the nutrient.

### Origin of the power-law growth–lag trade-off

Upon a sudden shift from P to NP, the pre-existing high level of *ϕ* suppresses flux from *n* → *g* and dissipates excess accumulation of *n*. This depletes cellular resources and delays the reversal of net flux. Because *ϕ* decays only on the timescale ~1*/µ*, flux reversal stalls until *ϕ* falls and *ζ* rises toward 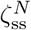 so that the growth rate *µ* can rise. This leads to a lag before growth can resume.

The metabolic model operates in at least three qualitatively distinct regimes, delineated by 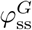, depending on how the system produces new enzymes *ζ* after the switch from P to NP. Here, we expand on the third scenario discussed in the main text, where 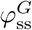 has intermediate values such that the value of 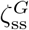 that maximizes pre-shift growth rate is zero (Fig. S5). We show below that the growth-lag trade-off in this regime is a power-law with exponent 1*/*(*ν* − 1).

Suppose 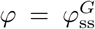 and 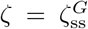 are the steady state values of the enzymes in P. At steady-state, *g* and *n* are found by solving

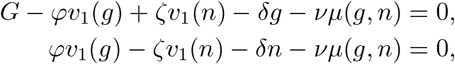

where *r* = 1, *v*_1_(*x*) = *x/*(*x*+*K*) (assume *K*_*g*_ = *K*_*n*_ = *K*), *µ*(*g, n*) = (*v*_2_(*g*)*v*_2_(*n*))^*ν*^ and *v*_2_(*x*) = *x/*(*x* + *K*_BM_).

We consider a parameter regime where, given 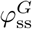, *µ*_P_ is maximized when 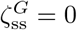 and consequently *ℓ* is infinite. We calculate how *µ*_P_ and *ℓ* decrease with 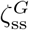 when 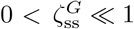.

The maximal growth rate when 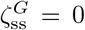 is found by solving the steady state equations

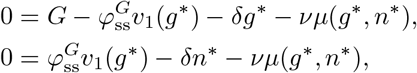

where *g*\*, *n*\* are the values at steady state in this scenario. Let Δ*g* = *g* − *g*\* and Δ*n* = *n* − *n*\* be the deviations from these values when 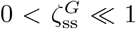. Δ*g* and Δ*n* are of order 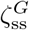. Expanding the steady state equations to first order when 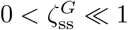, Δ*g* and Δ*n* satisfy

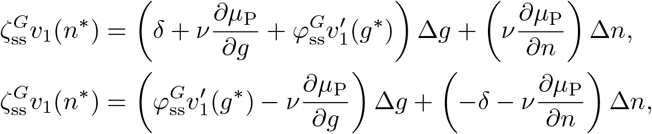

where the derivatives are evaluated at *g*\*, *n*\*. Since 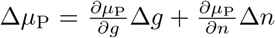 (to first order in 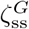), we solve Δ*g* and Δ*n* by solving the above pair of equations. This gives an expression for Δ*µ*_P_:

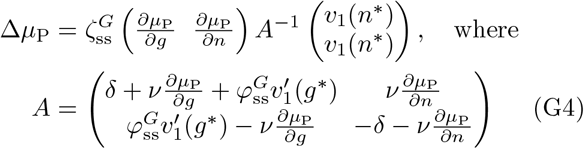

which we can write as 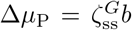, where *b* is a scalar obtained from the vector-matrix-vector product on the right hand side of the above equation.

After the switch to NP (starting from the steady state in P with 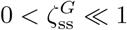), we have

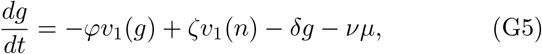

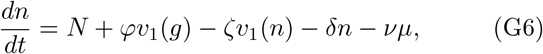

where

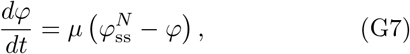

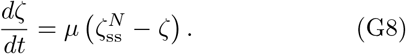

with initial conditions 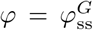 and 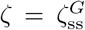. The approximations are valid soon after the switch. Dividing these two equations, we get 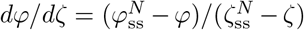. Integrating, we get

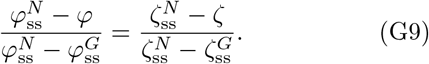

As discussed earlier, we work in the limit where after the switch, *g* depletes rapidly due to both the transformation from *g* to *n* catalyzed by *ϕ*, and due to loss at rate *δ*. In quantitative terms, the first and third terms on the rhs of (G5) contribute to the outflux of *g*.

During the lag phase, both *ϕ* and *ζ* do not change significantly from 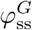 and 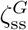 as biomass production (i.e., *µ*) is minimal in this phase. When 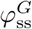 is large, *g* drops rapidly to near zero until it reaches a steady state. Since *v*_1_(*g*) ≈ *g* when *g* ≪ *K* and *v*_1_(*g*) ≈ 1 when *g* ≫ *K*, either *g* initially decreases exponentially or linearly with rate 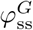 depending on the value of *g*\*. Moreover, when *g* ≪ 1, *µ* ≪ *v*_1_(*g*) as *µ* ~ *g*^*ν*^ and *v*_1_(*g*) ~ *g* for small *g* and *ν >* 1. That is, *dg/dt* ≈ − (*ϕ* + *δ*)*g* + *ζv*_1_(*n*). At steady state, we have (*ϕ* + *δ*)*g* ≈ *ζv*_1_(*n*).

Moreover, in steady state, from (G6) we see that all terms except *N* and *δn* are negligible so that *n* ≈ *N/δ*. Thus,

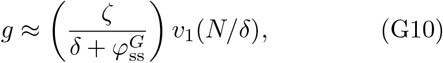

where have used (G9) to eliminate *ϕ* and ignored all terms second order in *ζ*. Plugging this expression for *g* into *µ* in (G8), we have

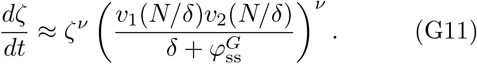

Integrating this from 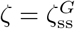 to ∞ (or some large value) for *ν >* 1, we get an estimate of lag *ℓ* as the time it takes for *ζ* to diverge:

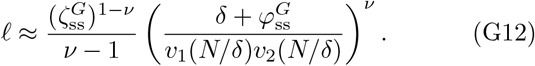

Plugging in 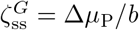, we have

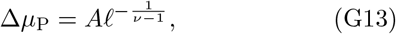

for some constant *A*.

**Fig. S5.**
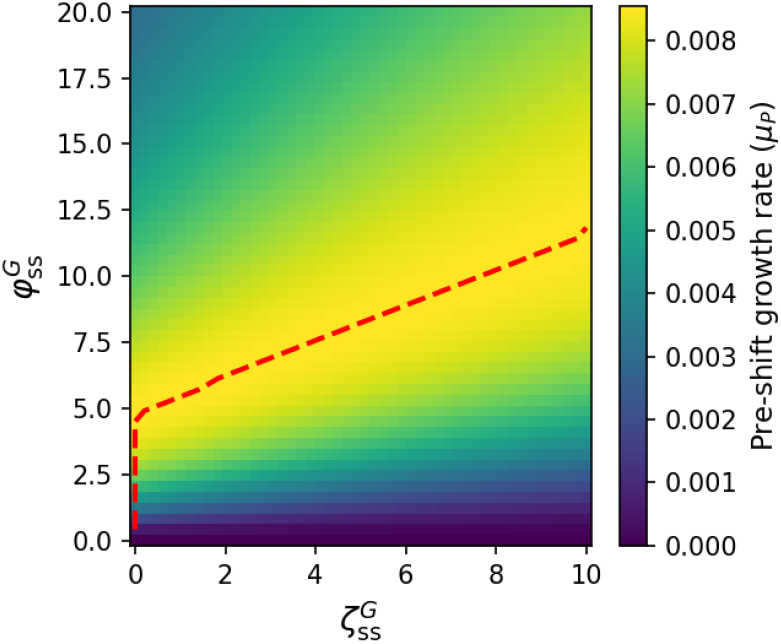
Heatmap of pre-shift growth rate *µ*_P_ for different 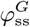 and 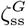 values. Other parameters in the model are fixed. Above a certain value of 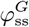, the optimal pre-shift growth rate is achieved at a non-zero 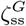 (dashed red curve).

## Notes

### Competing Interest Statement

The authors have declared no competing interest.

### Summary of Updates

Figure 3 and 4 revised. Added a new result of fitting theory to data.

